# Transcriptional profiling of non-injured nociceptors after spinal cord injury reveals diverse molecular changes

**DOI:** 10.1101/743567

**Authors:** Jessica R. Yasko, Isaac L. Moss, Richard E. Mains

## Abstract

Traumatic spinal cord injury (SCI) has devastating implications for patients, including a high predisposition for developing chronic pain distal to the site of injury. Chronic pain develops weeks to months after injury, consequently patients are treated after irreparable changes have occurred. Nociceptors are central to chronic pain; however, the diversity of this cellular population presents challenges to understanding mechanisms and attributing pain modalities to specific cell types. To begin to address how peripheral sensory neurons distal to the site of injury may contribute to the below-level pain reported by SCI patients, we examined SCI-induced changes in gene expression in lumbar dorsal root ganglia (DRG) below the site of injury. SCI was performed at the T10 vertebral level, with injury produced by a vessel clip with a closing pressure of 15g for 1 minute. Alterations in gene expression produce long-term sensory changes, therefore we were interested in studying SCI-induced transcripts before the onset of chronic pain, which may trigger changes in downstream signaling pathways and ultimately facilitate the transmission of pain. To examine changes in the nociceptor subpopulation in DRG distal to the site of injury, we retrograde labeled sensory neurons projecting to the hairy hindpaw skin with fluorescent dye and collected the corresponding lumbar (L2-L6) DRG 4 days post-injury. Following dissociation, labeled neurons were purified by fluorescence-activated cell sorting. RNA was extracted from sorted sensory neurons of naïve, sham, or SCI mice and sequenced. Transcript abundances validated that the desired population of nociceptors were isolated. Cross-comparisons to data sets from similar studies confirmed we were able to isolate our cells of interest and identify a unique pattern of gene expression within a subpopulation of neurons projecting to the hairy hindpaw skin. Differential gene expression analysis showed high expression levels and significant transcript changes 4 days post-injury in SCI cell populations relevant to the onset of chronic pain. Regulatory interrelationships predicted by pathway analysis implicated changes within the synaptogenesis signaling pathway as well as networks related to inflammatory signaling mechanisms, suggesting a role for synaptic plasticity and a correlation with pro-inflammatory signaling in the transition from acute to chronic pain.

**Contribution to the field:** Traumatic spinal cord injury (SCI) has devastating implications for patients, including a high predisposition for developing chronic pain. Much of the pain seems to emanate from tissues further away from the brain than the site of injury. Chronic pain develops weeks to months after injury, which means that patients are frequently treated only after enduring pain has developed. Nociceptors are the specialized sensory neurons central to chronic pain. We were interested in studying SCI-induced gene transcript (RNA) changes before the onset of chronic pain, in the hope of identifying mechanisms which could become therapeutic targets. Nociceptors below the site of spinal injury were isolated and their RNAs were sequenced. The results identified a unique pattern of gene expression in the subpopulation of nociceptors projecting to the relevant peripheral tissue. Particularly interesting were sets of genes crucial to synapse formation and maturation – the ability of neurons to talk to each other – and genes involved in inflammatory responses, since treatment of inflammation of nervous tissue could also be important for therapeutic approaches. It is evident that the transition from acute to chronic pain occurs in distinct steps that involve numerous signaling pathways, providing a host of potential new drug targets.

## Introduction

While spinal cord injury (SCI) is typically associated with loss of locomotor function, it can also result in chronic pain, affecting nearly 70% of patients with SCI (Finnerup et al. 2001). There are many categories of pain types affecting this population however, studies indicate that neuropathic pain below, or distal, to the level of injury is among the most common and difficult to treat (Defrin et al. 2001, Finnerup, Johannesen, Sindrup, Bach and Jensen 2001, Nees et al. 2016, Siddall and Loeser 2001, Yezierski 2005). Of those patients reporting below-level neuropathic pain, half described their pain as severe or excruciating, causing significant disability in patients already disabled from loss of motor function (Defrin, Ohry, Blumen and Urca 2001, Siddall et al. 2003). With few patients able to achieve complete relief with current treatment options, research has focused on mechanisms responsible for SCI pain at the site of injury, with the intention of treating the injury itself to prevent subsequent development of pain.

Considerable advances have been made in understanding changes within the spinal cord, including how spinally mediated alterations contribute to SCI-induced pain by increasing spinal cord excitability, and by establishing a variety of factors that impact how incoming sensory stimulation is processed (Bruce et al. 2002, Meisner et al. 2010, You et al. 2008). However, this approach has not translated into successful pain management. This may be attributed to an incomplete understanding of the differential functions of specific afferent subtypes in SCI, and how afferents distal to the site of injury become sensitized in patients with chronic below-level pain (Thakur et al. 2014). The sensory system receives inputs from multiple cell types, and peripheral cell bodies within the dorsal root ganglion (DRG) are important targets for assessing sensory function and pain (Usoskin et al. 2015). Persistent activity from injured and non-injured afferent fibers contributes to development and maintenance of chronic pain following SCI (Gold and Gebhart 2010).

Each sensory neuron has a unique pattern of gene expression that influences its modality-specific contribution to injury-induced pain (Le Pichon and Chesler 2014). To better understand the underlying pathophysiology of below-level pain following SCI, it is necessary to identify changes in cells impacted by the injury. The skin is heavily innervated by a broad range of nociceptors, and previous work has shown that SCI can impact the function of cutaneous nociceptors below the level of injury (Berta et al. 2017). This has been demonstrated by sustained spontaneous activity in peripheral terminals and in cell bodies of sensory neurons projecting to the skin after initial SCI (Bedi et al. 2010, Carlton et al. 2009, Ritter et al. 2015, Wu et al. 2013, Yang et al. 2014). Additional work has demonstrated that blockade of peripheral afferents into the central nervous system can effectively mitigate patient discomfort and chronic pain (Basbaum et al. 2009, Campbell et al. 1988, Gold and Gebhart 2010). These data support the idea that the mechanisms generating and maintaining prolonged pain reside within the peripheral nervous system.

In the present study, we identify specific transcriptional alterations in non-injured DRG distal to the site of injury. Using retrograde labeling from hairy hindpaw skin and flow cytometry, we isolated a nociceptor population projecting to sites distal to the spinal injury, free of surrounding neuronal and glial cells. This enabled identification of novel cutaneous nociceptor genes and predicted pathways not discernible by whole DRG tissue analyses.

## Methods

### Animals

Experiments were conducted with adult (8-12 week) female C57BL/6J mice (Jackson Laboratory, Bangor ME). Several chronic pain conditions have a higher prevalence in females, and numerous studies have reported higher pain prevalence in the SCI population among female patients (Cardenas et al. 2004). Women also report greater frequency, severity, and longer lasting pain, as well as neuropathic pain below the level of injury, in comparison to men (Cardenas, Bryce, Shem, Richards and Elhefni 2004). The majority of research examines SCI in male rodents and this study will add to what is known in the literature by focusing on female mice (Cardenas, Bryce, Shem, Richards and Elhefni 2004). Naïve animals were group housed; sham and spinal cord injured animals were individually caged. All animals were maintained on a 12:12-h light-dark cycle with a temperature-controlled environment, and given food and water ad libitum. All treatments and testing were approved by the University of Connecticut Health Center Institutional Animal Care and Use Committee.

### Spinal cord injury (SCI) procedure

Animals were anesthetized by inhalation of isoflurane and a 1.0-cm dorsal midline skin sterile incision was made over T8-T11, as per Ma *et al*. (Ma et al. 2001). Connective and muscle tissue were removed to expose the bone from T9-T10, and a laminectomy was performed at the T10 vertebral level. Spinal cord injury was produced by compression of the vertical plane of the spinal cord using a vessel clip with a closing pressure of 15g (WPI, Sarasota, FL) for 1.0 minute, exerting pressure from side to side on the spinal cord. Sudden impact is produced by the rapid release of the vessel clip (Tator 2008). This injury is analogous to the majority of lesions in humans, as the model constitutes both contusion and compression (Marques et al. 2014). After removal of the clip a hemorrhagic ring at the site of compression is present. The wound is closed with coated vicryl absorbable sutures (Ethicon, Somerville, NJ). Mice were allowed to recover in warm cages for 24hr. All animals were administered antibiotics once immediately following surgery (5mg/kg gentamicin), as well as subcutaneous saline for 4 days following surgery, without analgesics. Sham control mice received the same treatment excluding the vessel clip. Manual bladder expression on SCI mice was performed twice daily until mice were sacrificed. Mortality was less than 10% and typically occurred during the laminectomy due to excessive blood loss, or when postinjury weight loss required the animal to be sacrificed. Spinal cords collected at 4 days post-injury were most notably characterized by minimal cavitation and scar tissue that progressively diminished with increasing distance distal to the site of injury (~L1-2 vertebral level), analogous to previous studies that have characterized this injury model (Joshi and Fehlings 2002, Marques, de Almeida, Mostacada and Martinez 2014).

## Behavioral tests

### Tail-flick test for thermal sensitivity

Mice were acclimated in 50mL tubes for 2 days prior to testing, 20 minutes per day. On testing days, mice were left in their home cage to acclimate to the test room for 30 minutes before testing (Bannon and Malmberg 2007). Latency to respond to thermal stimuli was measured by dipping the distal 1.5cm of the tail into a 50°C water bath (Ramabadran et al. 1989). The tail was removed from the water upon response, or after 15 seconds to prevent tissue damage. The stimulus was conducted 3 times, at 20 second intervals or less (Zhou et al. 2014). The first response was dropped, and the average latency to respond from two trials was used for analysis. Video recording and VLC software were used to determine tail-flick responses in milliseconds. Mice were tested for thermal sensitivity 1 day prior to surgery for baseline response thresholds, and at days 1, 3, 5, and 7 post-surgery. N=6 per group for each time point.

### Mechanical sensitivity

To assess mechanical sensitivity, mice were confined in clear plastic containers placed on an elevated wire mesh platform. Prior to testing, mice were acclimated to the apparatus for 60 minutes. Mechanical reactivity was assessed on the plantar surface of the hind paw using a series of calibrated von Frey filaments according to the up-down method as described (Dixon 1980), and 50% response thresholds were compared across all conditions. Both hindpaws were tested for mechanical sensitivity, and collapsed across each group of mice per condition. Mice were tested for mechanical hypersensitivity 1 day prior to surgery for baseline response thresholds, and at days 1, 3, 5, and 7 post-surgery. N=6 per group for each time point.

### Open Field Test

The open field test was conducted using a 16”x16” open-field container subdivided by infrared beams to track movement (San Diego Instruments, San Diego, CA). Data were acquired using the manufacturer’s tracking software, which records ambulation movements based on beam breaks as well as central vs. peripheral beam break counts. All mice were placed in the same corner of the box before testing and allowed to freely explore for 10 minutes. Mice were tested 1 day prior to surgery for baseline locomotor behavior, and at days 1, 3, 5, and 7 post-surgery. Spinally injured mice were tested 1 day prior to surgery and 1 day post-surgery. Naive N=4 for each time point; Sham N=3-10 for each time point (within group design, tissue was collected for ELISAs for corresponding time points); N=4 SCI day 1.

### Cuprizone treated mice

Female C57BL/6 mice (6-10 weeks old) were fed powdered milled chow mixed to contain a final concentration of 0.2% bis (cyclohexanone) oxaldihydrazone (cuprizone; Sigma-Aldrich, St. Louis, MO), with food and water available *ad libitum*. Each mouse received approximately 5g of chow per day, fresh cuprizone containing chow was prepared every 7 days. Cuprizone feeding was maintained for 35 days, and tissue was collected for protein analyses on day 35. Digitized, non-overlapping electron micrographs of the corpus callosum were analyzed for unmyelinated axon frequency and g-ratios to assess effectiveness of cuprizone treatment (Wasko et al. 2019). N=5.

### Cytokine ELISAs

Spinal cord segments at the level of laminectomy (T8-T11) were collected from naïve, sham, SCI, or cuprizone treated mice immediately following perfusion with ice cold 0.9% NaCl. Spinal cord segments were homogenized in ice-cold buffer containing 20mM TES, pH 7.4, 10mM mannitol, 0.3mg/mL phenylmethylsulfonyl fluoride, 2μg/mL leupeptin, 2μg/mL pepstatin, 2 μg/mL benzamidine, 16μg/mL benzamidine, and 50μg/mL lima bean trypsin inhibitor at a concentration of 0.1g tissue per 1mL buffer (Mains et al. 2018). Homogenates were freeze-thawed three times, centrifuged (20min, 17,400g), and supernatants were collected. Approximately 60 g of protein per sample was used for each ELISA. The ELISA assays were performed according to the manufacturer’s instructions (R&D systems mouse duo-sets IL-10, IL-6, IL-1β, TNF-α, completed with Ancillary Reagent Kit 2 Minneapolis, MN). The sample absorbance was read with an ELISA plate reader at 450nm; readings were also taken at 570nm to subtract optical background. The concentration was determined based on a standard curve. All results were normalized to amount of protein added per sample and graphed as pg/mg. Naïve, 1d Sham, 4d Sham, and 1d SCI N=4 mice; 5d and 7d Sham conditions N=3 mice.

### Backlabeling procedure

To backlabel DRG L2-L6 projecting to the hairy hindpaw skin, mice were anesthetized with isoflurane. 0.3% wheat germ agglutinin conjugated to an AF-488 dye (WGA-488, Thermo Fisher, Waltham, MA) in sterile PBS was injected into the sural, common peroneal, and saphenous nerve skin territories for retrograde labeling of DRG neurons (Berta, Perrin, Pertin, Tonello, Liu, Chamessian, Kato, Ji and Decosterd 2017, da Silva Serra et al. 2016). A total of 6μL of WGA-488 was injected 2 days prior to surgery by three 2μL injections in the lateral zones of each hindpaw (2μL per nerve territory) using a 10μL Hamilton Syringe and 30G needle. This was performed on both hindpaws of each mouse. This technique does not cause significant injury to the sensory afferents being studied.

### Primary DRG neuron dissociation

Mice were anesthetized 4 days post-surgery with an intraperitoneal injection of ketamine (100 mg/kg) plus xylazine (10 mg/kg) and perfused with ice cold 0.9% NaCl. A laminectomy was performed and L2-L6 DRG from both sides of the spinal column were collected into cold HBSS (KCl 5.4mM, NaCl 137mM, Glucose 5.6mM, Hepes 20mM, pH 7.35 NaOH), after which the mice were sacrificed by decapitation. Sensory neuron dissociation was performed as described (Malin et al. 2007). Briefly, following collection, tissue was treated with 60U papain (Worthington), 1mg of cysteine, and 6μL of NaHCO_3_ in 1.5mL HBSS at 37°C for 10 min. Tissue was then treated with 12mg collagenase II (Worthington, Lakewood, NJ) and 14mg dispase (Roche, Basel, Switzerland) in 3mL HBSS at 37°C for 20 min, washed, and triturated with fire polished glass Pasteur pipettes in 1mL of DMEM (Gibco Thermo Fisher Scientific, Waltham, MA) supplemented with FBS (Hyclone, Logan, UT) and pen/strep (Gibco). The cell suspension was pelleted (1 min, 80g), DMEM was removed, and cells were re-suspended in a modified solution (Citri et al. 2011) containing 140mM NaCl, 5mM KCl, 10mM Hepes, 10mM glucose, 0.1% Bovine Serum Albumin, pH 7.4. After re-suspension, cells were strained through a 70μm cell strainer and placed on ice in the modified solution until fluorescence activated cell sorting (FACS).

### Imaging Flow Cytometry

Single cell suspensions of cells isolated from *in situ* WGA-488 labeled DRG were live-stained using Hoechst 33342 (10 g/mL, Thermo Fisher) and propidium iodide (PI) (1 g/mL), and analyzed on an Amnis ImageStreamX Mark II imaging flow cytometer (Luminex Co., Austin, TX). Fluorescent cell images were captured using a 60x objective lens with excitation from a 405nm laser at 20mW power and a 488nm laser at 200mW power. Images of in-focus nucleated WGA AF488-positive cells were identified and electronically gated using IDEAS software (Amnis, v6.2.183, Seattle, WA).

### Flow cytometry and cell sorting

Neurons labeled with WGA-488 dye *in situ* in the DRG were purified by fluorescence activated cell sorting 4 days post-surgery. Following primary dissociation of DRGs L2-L6, single cell suspensions were analyzed and sorted using a BD FACS Aria II cell sorter (Becton Dickinson) set up with a 130μm nozzle at 12 PSI in order to gently isolate cells between 10 and 30μm. Single live neurons were defined by electronic gating in FACS DIVA software (BD, ver. 8.01) using forward and side angle light scatter, omission of propidium iodide (PI, 1 g/mL), and AF488 fluorescence. All fluorescence gates were confirmed using fluorescence minus one controls (e.g.: a sample of cells from unlabeled DRG was used to gate for AF488 positive cells and a sample of cells not treated with PI was used to set the live cell gate). WGA-488 positive cells were sorted directly into lysis buffer (NucleoSpin RNA XS Kit, Machery-Nagel, Bethlehem, PA) and immediately placed on dry ice until RNA extraction.

### RNA extraction and RNA sequencing

RNA from FACS sorted cells was isolated using NucleoSpin RNA XS Kit, including a DNA digestion step but without carrier RNA step. Before library preparation RNA quality and integrity was tested for each sample using the Agilent High Sensitivity RNA Screen Tape on the Agilent Tapestation 2200 (Agilent Technologies, Santa Clara, CA). RNA with RIN values ≥6.7 (minimum 6.7, maximum 9.9, average 7.3) was further processed for RNA sequencing. Library preparation was performed using the Illumina sequencing kit for high output 75-cycles for 25-30M total single end reads per sample. DESeq2 analyses (https://bioconductor.org/packages/release/bioc/html/DESeq2.html) of differential expression were performed, and outliers beyond 30-50% of the mean for each group of animals were eliminated (Conesa et al. 2016, Labaj and Kreil 2016, Love et al. 2014, Wu and Wu 2016).

### Pathway Analysis

Data was analyzed by Ingenuity Pathway Analysis (IPA; Qiagen, Germantown, MD). An overlap of significance for DEseq2 comparisons plus an RPKM cutoff >10 were required for transcripts to be included for IPA analysis. We analyzed 125 transcripts for comparisons between SCI and naïve groups, and 560 transcripts for comparisons between SCI and sham groups.

## qPCR validation

### Pre-amplification of cDNA for Gene Expression

cDNA was generated from RNA samples from FACS sorted cells with the iScript Reverse Transcription Supermix (#1708840 Bio-Rad, Hercules, CA), N=6 per condition. Target-specific preamplification was performed on cDNA generated from RNA samples using SsoAdvanced PreAmp Supermix (#1725160 Bio-Rad) containing Sso7d fusion polymerase. Briefly, 20 L of cDNA was preamplified in a total volume of 50 L containing 25 L of 2x SsoAdvanced PreAmp Supermix and 21 primer pairs, 50nM of each primer. Preamplification was performed at 95°C for 3 min followed by 12 cycles of amplification at 95°C for 15 seconds and 58°C for 4 min. Samples were moved directly to ice and stored at −80°C. Preamplified cDNA were diluted 1:5 with H_2_O.

### qPCR

Following cDNA synthesis and preamplification, qPCR was performed using the primers listed in **Suppl. 6**. All primers had calculated melt temperatures of 59.5-63.5C, and all products were 111-143 bp in length, as verified by agarose gel electrophoresis. qPCR was performed at 95°C, 2 min; 95°C, 10 seconds; 55 °C, 15 seconds; and 72°C, 40 seconds, repeating the second through fourth steps for a total of 40 cycles in a Bio-Rad CFX Connect Optics Module machine. iQ SYBR Green Supermix (#1708882 Bio-Rad) was used for linear detection of qPCR results. Hypoxanthine phosphoribosyltransferase (Hprt) was used as the most constant normalizer transcript, based on the RPKM data (Klenke et al. 2016, Lima et al. 2016).

### Statistical analyses

Differences between groups were compared using Student’s t-test or ANOVA, followed by Tukey’s posttest, Bonferroni’s multiple comparisons test, or by unpaired Student’s t-test. P-values <0.05 were considered statistically significant. Statistics on PCR data were conducted using delta CT values. Heat maps were generated by Microsoft software (Excel), hierarchical clustering was generated by Gene Cluster 3.0 and visualized using Java TreeView (Baek et al. 2017, de Hoon et al. 2004). All other data were plotted using Prism 6 (GraphPad Software, San Diego, CA). R studio was utilized for differential expression analysis; Prism software was used for all other statistical tests.

## Results

### Characterization of behavioral and inflammatory phenotypes of sham mice

To determine an optimal time point to observe transcriptional changes contributing to the transition from acute to chronic pain, we tested behavioral differences between naïve and sham operated mice. This was to ensure that changes within the DRG were due to injury to the spinal cord itself, not to the laminectomy performed in both injured and sham mice. Specifically, we tested naïve and sham mice for open field locomotor differences 1, 3, 5, and 7 days post-surgery (**Fig.1A1**). Naïve and sham mice did not differ significantly at any of the time points tested, including as early as 1 day post-surgery. To ensure that SCI mice (T10 compression-clip injury) did exhibit behavioral differences and locomotor deficits following injury, we compared SCI mice to naïve and sham mice for open field behavior 1 day post-injury (**Fig.1A2**). SCI mice exhibited substantially decreased total ambulation (one-way ANOVA, p=0.0059, Tukey’s multiple comparisons test, naïve **p<0.005, sham *p<0.05) following injury, as expected, although time spent in the periphery did not differ (**Suppl.1A**). Additional tests for mechanical (von Frey) and thermal (hot water tail-flick) hypersensitivity showed that naïve and sham conditions did not differ significantly at any time point (**Fig.1B**). We did not examine SCI mice for thermal or mechanical sensitivity, as paralysis below the level of injury prohibited below-level sensitivity testing and previous work using the clip-compression model has not demonstrated significant changes in above-level sensitivity following SCI (Bruce, Oatway and Weaver 2002). The development of chronic pain at later time points following clip-compression models of SCI (in addition to several other models) has already been well characterized. Therefore we focused on early time points after injury, to identify contributions to the onset of pain, rather than changes after chronic pain has already developed (Bruce, Oatway and Weaver 2002, Gaudet et al. 2017, Nakae et al. 2011).

**Figure 1.**
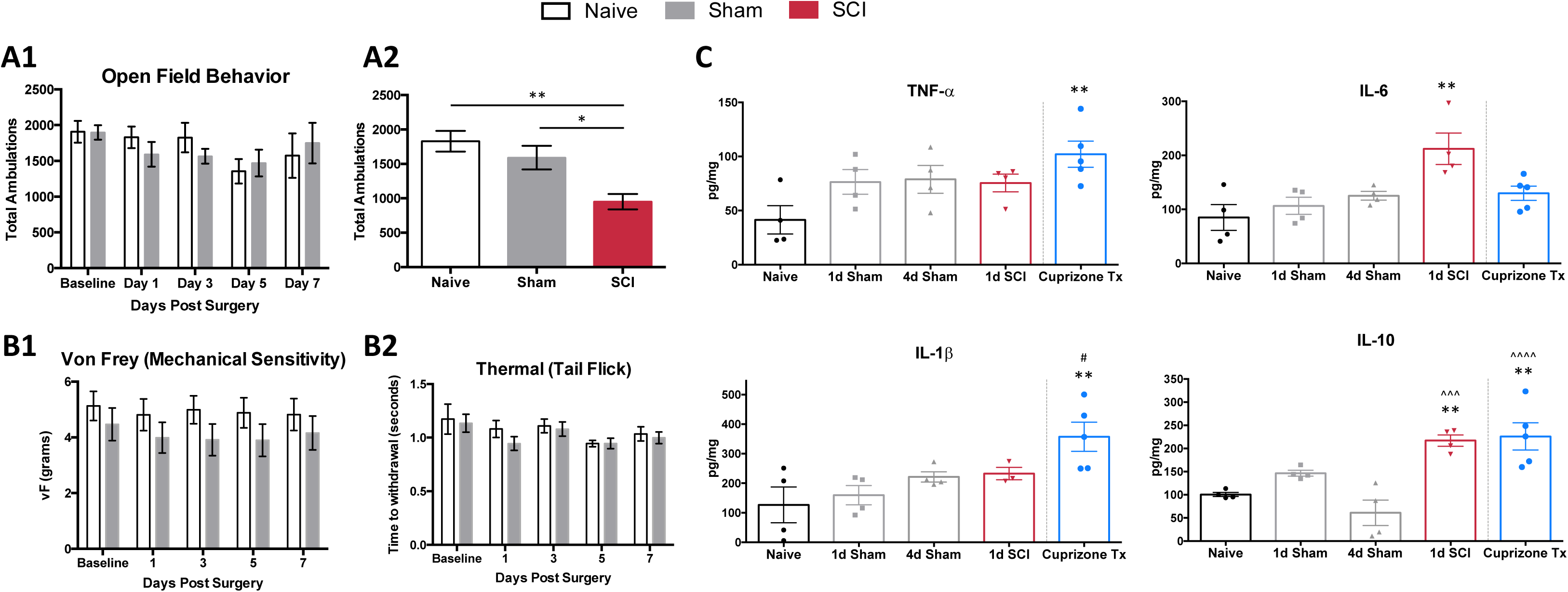
Behavioral responses and cytokines. (A1) Open field behavior (10 minute trials) in naïve or sham mice 0,1,3,5 and 7 days post-surgery does not differ significantly at any time point in total ambulation, N=3-10 sham, 4 naïve. (A2) Total ambulation differs significantly 1 day post-SCI in both SCI vs. naïve (one-way ANOVA, p=0.0059, Tukey’s multiple comparisons test **p<0.005) and SCI vs. sham (*p<0.05) mice. (B1) Mechanical and (B2) thermal sensitivity do not differ significantly 0,1,3,5, and 7 days post-surgery in naïve and sham mice, N=6 each. (C) Cytokine ELISAs on spinal cord segments at the level of laminectomy (T8-T11) show no significant differences between naïve and sham mice 1 or 4 days post-surgery. 1-day post-operation, SCI mice have significantly increased levels of IL-6 and IL-10 compared to naïve controls (one-way ANOVA, Bonferroni’s multiple comparisons test; *p<0.05, **p<0.005, respectively) and significantly increased levels of IL-10 in comparison to 4d sham condition (one-way ANOVA, Bonferroni’s multiple comparisons test; ***p<0.001). Cuprizone treated mice as positive control, with significantly increased TNF-α, IL-1β, and IL-10 compared to naïve controls (one-way ANOVA, Bonferroni’s multiple comparisons test; **p<0.005, **p<0.005, ***p<0.001, respectively) after 5 weeks of cuprizone treatment and significant increases in IL-1β at the 1d sham timepoint, and IL-10 at the 4d sham timepoint (one-way ANOVA, Bonferroni’s multiple comparisons test; *p<0.05, ****p<0.0001, respectively). * Represents comparisons to naïve conditions, # represents comparisons to 1d sham conditions, ^ represents comparisons to 4d sham conditions.

Previous work has highlighted the importance of inflammatory cytokines within the CNS to facilitate the transduction of noxious stimuli in neuropathic pain (Cook et al. 2018). Thus, we examined post-surgical changes in inflammation, using ELISAs to analyze common cytokine markers to ensure that sham mice did not differ significantly from naïve mice following the removal of bone and muscle. We did not observe significant differences in inflammatory cytokine levels (TNF-α, IL-6, IL-1β, IL-10) in extracts of spinal cord segments (T8-T11) in naïve or sham mice at 1, 4, 5, and 7 days post-surgery (**Fig.1C**, **Suppl.1B**). Spinal cords from SCI mice had significantly increased levels of IL-6 and IL-10 in comparison to naïve controls (one-way ANOVA, Bonferroni’s multiple comparisons test; *p<0.05, **p<0.005, respectively). When compared to sham conditions, spinal cords from SCI mice had significantly increased levels of IL-10 in comparison to 4d, 5d, and 7d sham conditions (one-way ANOVA, Bonferroni’s multiple comparisons test; ***p<0.001, **p<0.005, **p<0.005, respectively. As a positive control, we tested cuprizone-treated mice (a model of multiple sclerosis), which are known to secrete proinflammatory cytokines at the end of 5 weeks of treatment (Mukhamedshina et al. 2017). Cuprizone-treated mice showed expected increases in levels of TNF-α, IL-1β, and IL-10 in comparison to naïve controls (one-way ANOVA, Bonferroni’s multiple comparisons test; *p<0.05, **p<0.005, **p<0.005, respectively) (Schmitz and Chew 2008). Similarly, in comparison to sham mice, spinal cord extracts from cuprizone-treated mice had significant increases in IL-1β at the 1d sham timepoint, and IL-10 at the 4d, 5d, and 7d sham timepoints (one-way ANOVA, Bonferroni’s multiple comparisons test; *p<0.05, ****p<0.0001, **p<0.005, ***p<0.001, respectively). Behavioral testing and cytokine analyses did not reveal differences between naïve and sham mice at any time point.

### Confirmation of cell population specific labeling of cutaneous nociceptors

Because the laminectomy did not significantly affect behavior or inflammatory responses in sham mice, we determined an optimal time point to study the transition of acute to chronic pain based upon established characteristics of nociceptors following SCI. The inflammatory cytokine assays demonstrated that SCI causes major inflammation compared to sham controls at 4d after injury (**Fig.1C**, **Suppl.1B**). In addition, previous studies have documented onset of spontaneous activity in nociceptors distal to the site of SCI as early as 3 days post-injury; increased activity persisted for at least 8 months (Bedi, Yang, Crook, Du, Wu, Fishman, Grill, Carlton and Walters 2010). Thus, it is possible that the transition to chronic pain begins around 3 days after SCI, and this transition is not due to laminectomy, as nociceptors isolated from sham mice show no significant changes in spontaneous activity (Bedi, Yang, Crook, Du, Wu, Fishman, Grill, Carlton and Walters 2010, Yang, Wu, Hadden, Odem, Zuo, Crook, Frost and Walters 2014). Subsequently, we chose 4 days post-injury to assess transcriptional changes in nociceptors below the level of injury.

To perform transcriptional profiling on nociceptors that project to the cutaneous skin after injury, we injected wheat germ agglutinin conjugated to an AF-488 dye (WGA-488) into the sural, common peroneal, and saphenous nerve skin territories for retrograde labeling of DRG neurons. Next, we performed compression-clip SCI or sham surgeries at the T10 vertebral level 2 days post-WGA injection, and collected L2-L6 DRG for dissociation 4 days post-injury (**Fig.2A**). Based on the vertebral level at which we performed SCI, the DRG collected were located below the level of injury, thus our analysis was comprised of non-injured nociceptors that do not project directly to the lesion site. Flow cytometry confirmed that our cell population of interest (cutaneous nociceptors) was positively labeled with WGA-488 (**Fig.2B**). We also observed non-labeled (WGA-488) cells and dead cells (propidium iodide, PI+), to be excluded from cell sorting and analysis (**Fig. 2B**). As expected, a significant number of viable small nociceptor cells were excluded in this analytical approach, to avoid including RNAs from non-nociceptor cells or attached fragments of dead cells (examples in **Fig. 2C**) (Lopes et al. 2017, Megat et al. 2019, Thakur, Crow, Richards, Davey, Levine, Kelleher, Agley, Denk, Harridge and McMahon 2014).

**Figure 2.**
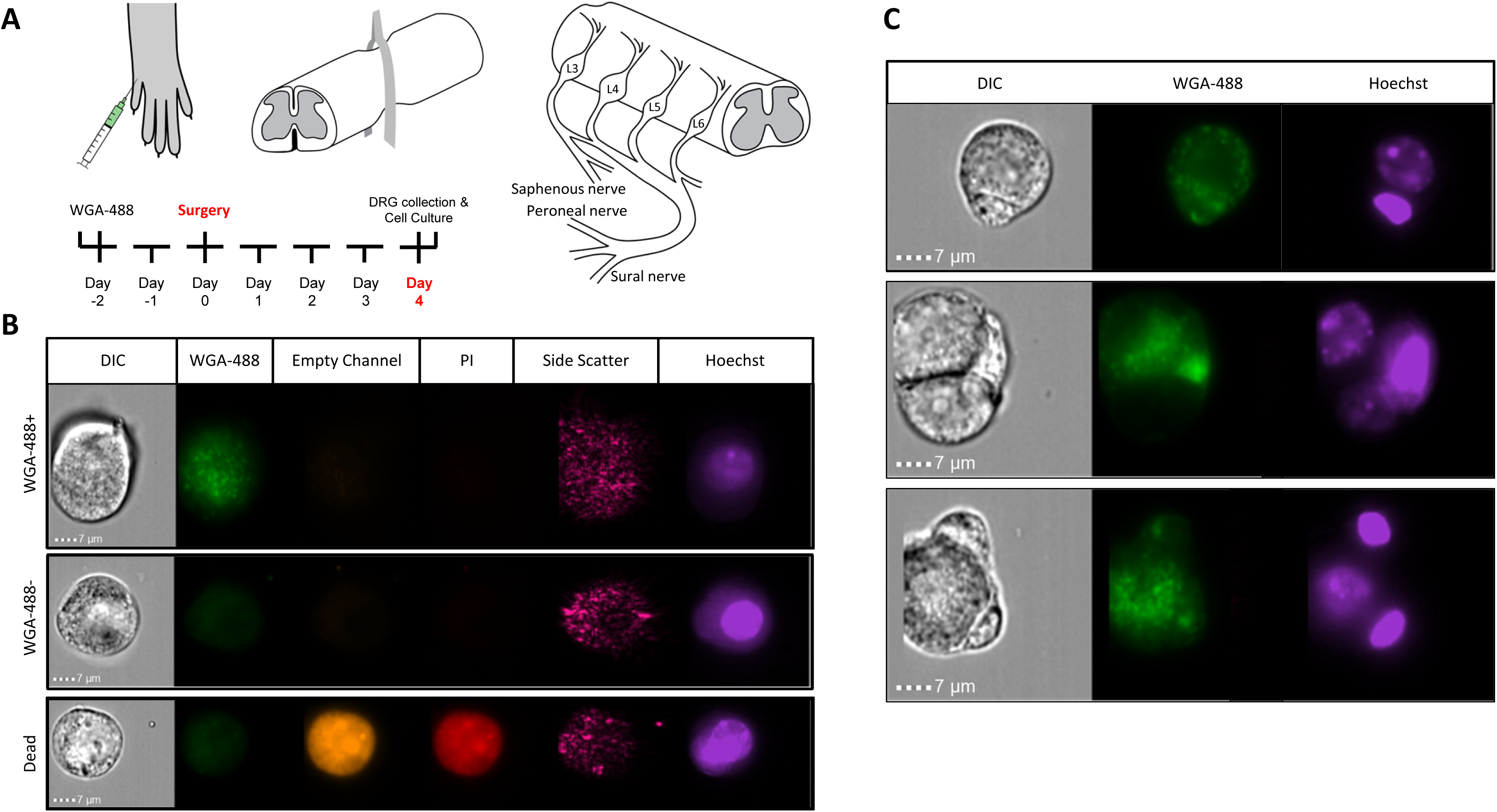
Identifying sensory neurons. (A) Diagram of WGA-488 injection under the hairy hindpaw skin of a mouse, how the spinal cord injury was conducted with a vessel clip, and projections to the corresponding DRG. (B) Fluorescent cell images taken by Amnis ImageStreamX Mark II imaging flow cytometer (Luminex Co.) using a 60x objective lens with excitation from a 405nm laser and a 488nm laser to identify propidium iodide (PI; red) positive (dead) cells and WGA (green) labeled cells, respectively. (C) Sample images of rejected cells because of attachment to non-nociceptor cells. Similarly, nociceptors with attached PI+ debris were eliminated.

### FACS purification of DRG nociceptors projecting to the cutaneous hind paw

We performed fluorescence-activated cells sorting (FACS) purification of nociceptor populations from naïve, sham, and SCI adult (8-12 week old) female mice (n=5 per condition). We pooled DRGs from lumbar regions L2-L6 on either side of the spinal column to ensure all DRGs isolated would have projections to the hairy hind paw skin. DRG cells were enzymatically dissociated and subjected to flow cytometry, to gently isolate positively labeled cells between 10 and 30μm; propidium iodide staining was used to identify dead cells. All conditions were gated on DRG from naïve mice that did not receive WGA-488 injections, enabling purification of positively labeled cells (**Fig.3A**). Analysis of our flow cytometry data shows that we were successful in retrogradely labeling DRG neurons projecting to hairy hind paw skin. Many positively labeled neurons were part of cell aggregates, limiting retrieval of the single cell population of interest to ~2% of all dissociated cells per animal (**Fig.3B**). This percentage amounted to approximately 3,000 cells per mouse, a suitable representation of this cell population in agreement with previous studies (Berta, Perrin, Pertin, Tonello, Liu, Chamessian, Kato, Ji and Decosterd 2017, Goswami et al. 2014, Thakur, Crow, Richards, Davey, Levine, Kelleher, Agley, Denk, Harridge and McMahon 2014, Usoskin, Furlan, Islam, Abdo, Lonnerberg, Lou, Hjerling-Leffler, Haeggstrom, Kharchenko, Kharchenko, Linnarsson and Ernfors 2015). DRG populations were sorted directly into lysis buffer and placed on dry ice to preserve transcriptional profiles at the time of isolation. RNA quality was tested using the Agilent TapeStation; a representative image of an RNA sample following FACS purification is shown (**Fig.3C**).

**Figure 3.**
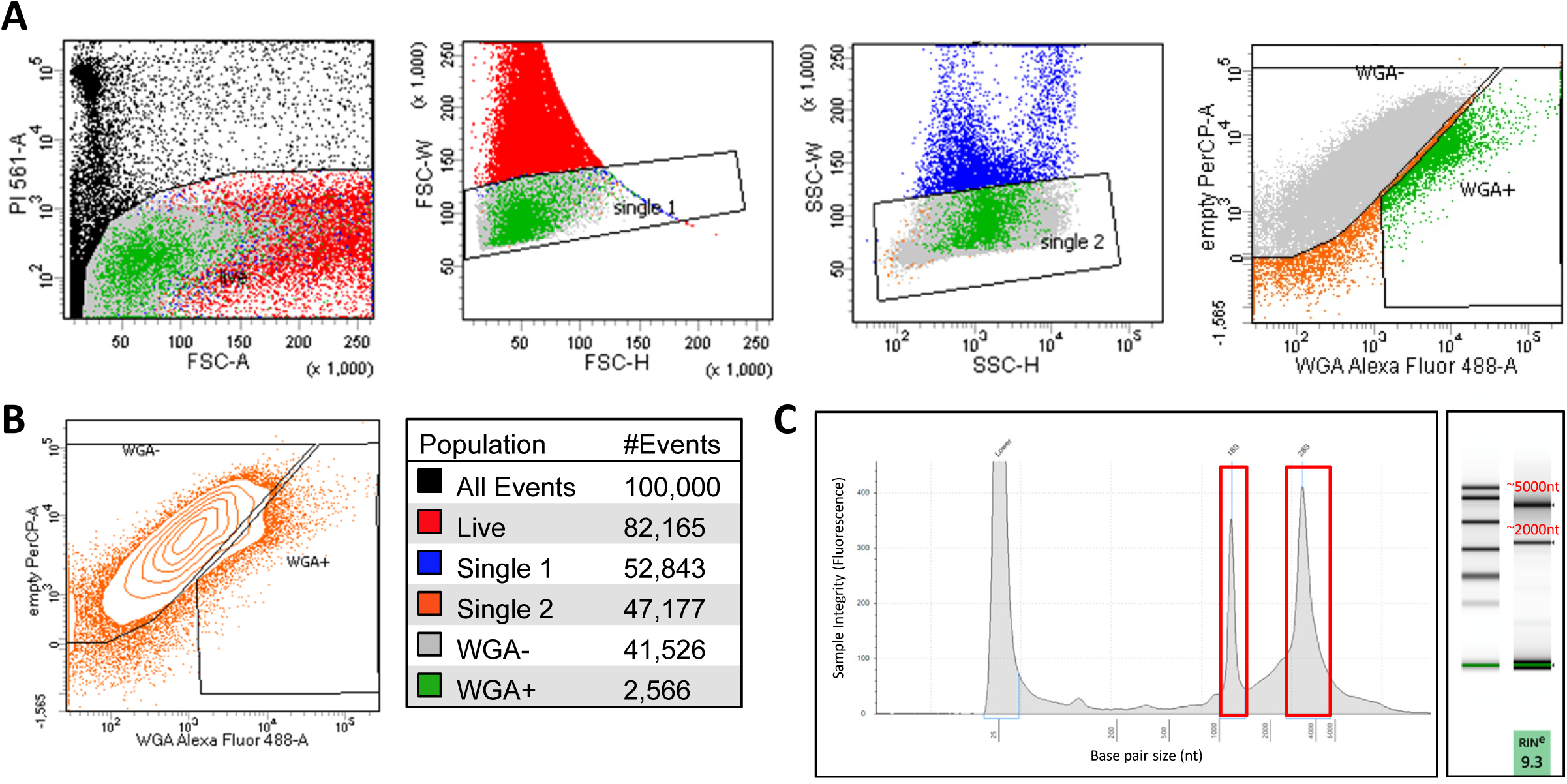
Isolating sensory neurons. (A) FACS purification of cultured DRG cells based on propidium iodide (PI) staining (dead) single cell populations by forward scatter (FSC), single cell populations by side scatter (SSC), and cells fluorescently labeled with WGA-488 (WGA+). (B) Contour plot of single WGA+ cells that were sorted directly into lysis buffer for RNA extraction and color scheme for the gated populations, each condition N=5; ~3000 cells. (C) Representative image of RIN values run on an Agilent TapeStation before RNAseq.

### Major characteristics of somatosensory mediators in the purified neuron population

We used the RNA sequencing data to evaluate the neuronal population that had been isolated. Scn10a, which encodes Na_v_1.8, is present in 80-90% of nociceptors and Trpv1 serves as a marker for the peptidergic population of nociceptors (Basbaum, Bautista, Scherrer and Julius 2009, Harriott and Gold 2009, Wu, Yang, Crook, O’Neil and Walters 2013). RPKM values of 1000 for Scn10a and 400 for Trpv1 confirm that FACS purified cells express high levels of these nociceptor markers **Fig.4A**). The low RPKM values observed for Parvalbumin (Pvalb), a glial transcript also found in large diameter proprioceptors and Aβ neurons, and Gfap (glial fibrillary acidic protein), another marker for glial cells, confirmed the absence of the non-nociceptive sensory neurons responsible for touch and proprioception (Aβ and Aδ neurons) and the absence of satellite glial cells from our purified nociceptor population (**Fig.4B**) (Huang et al. 2013, Le Pichon and Chesler 2014). Previous studies have confirmed that intact DRG as well as unsorted dissociated DRG yield much higher levels of nonneuronal markers (Thakur, Crow, Richards, Davey, Levine, Kelleher, Agley, Denk, Harridge and McMahon 2014). This highlights the importance of excluding cell aggregates during cell sorting (**Fig.3**) to avoid the analysis of transcripts from cells outside of the target population of nociceptors.

**Figure 4.**
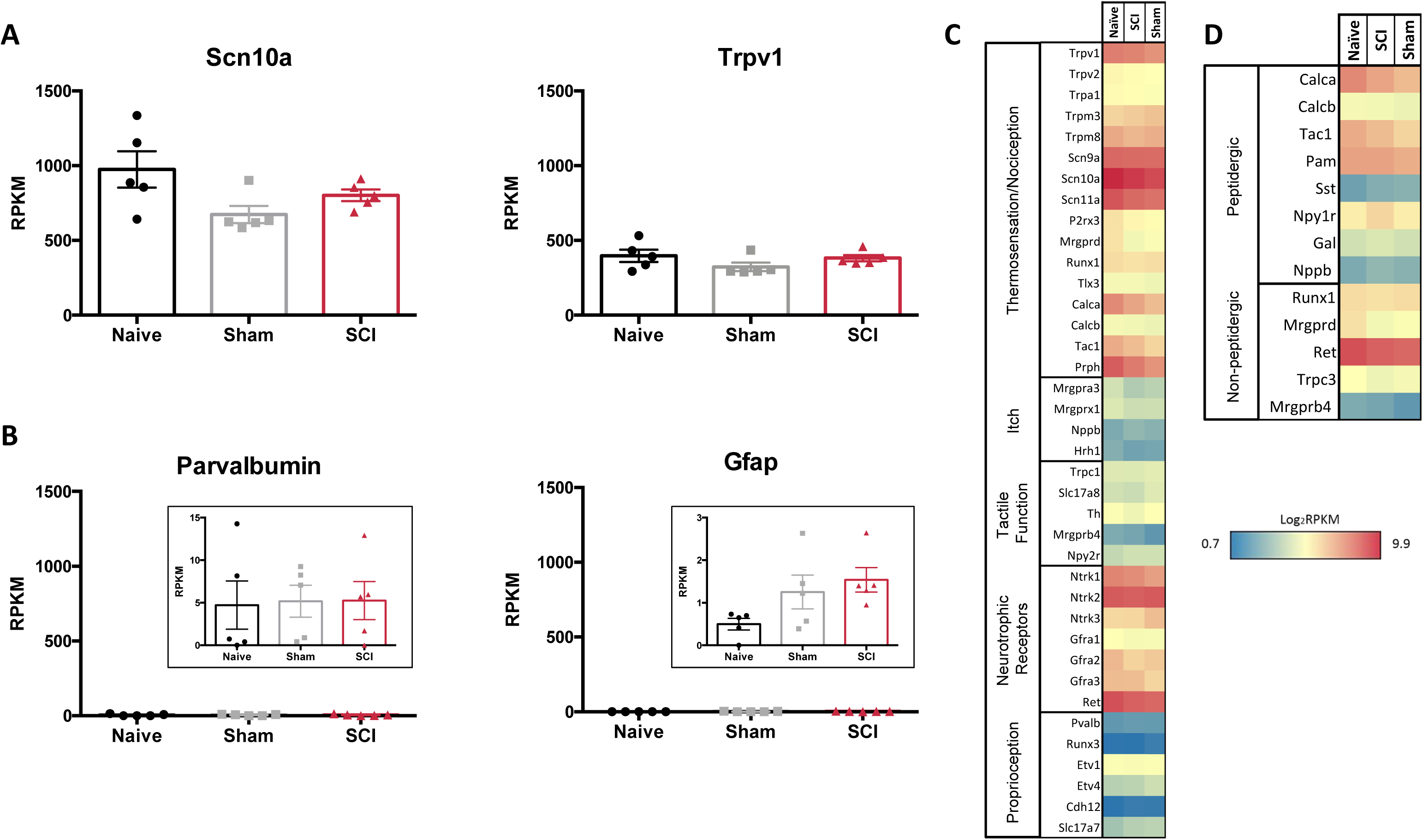
RNAseq demonstrates specificity of cell isolation. (A) Average RPKM values from RNAseq data set from genes that are known to be highly expressed in the cell population of interest or (B) expected to be expressed in other cell populations outside of the one being studied; larger myelinated neurons or satellite glial cells, respectively, N=5 per condition. (C) Heatmap of functional somatosensory mediators within the isolated cell population of interest. Genes were grouped based on roles established in the literature: thermosensation and nociception, itch, tactile function, and neurotrophic factors. (D) Heatmap of peptidergic and non-peptidergic markers. Graph: log_2_ of average RPKM values within each condition, RPKM >1, N=5 per condition.

We next analyzed gene expression patterns for known functional mediators of somatosensation (Chiu et al. 2014, Le Pichon and Chesler 2014, Usoskin, Furlan, Islam, Abdo, Lonnerberg, Lou, Hjerling-Leffler, Haeggstrom, Kharchenko, Kharchenko, Linnarsson and Ernfors 2015). The purified cutaneous nociceptors displayed high expression levels of genes involved in thermosensation and nociception, such as specific Trp channels (notably Trpv1), sodium channels (Scn9a, 10a, 11a) and Prph (peripherin) (**Fig.4C**). Markers for non-peptidergic nociceptors were abundant, such as Mrgprd (Mas-Related G-Protein Coupled Receptor Member D), Runx1 (Runt related transcription factor 1), Ret (Ret proto-oncogene), and Trpc3 (transient receptor potential cation channel C3). As expected, transcripts enriched in peptidergic nociceptors were present, such as Calca and Calcb (Calcitonin Related Polypeptides) and Tac1 (the tachykinin precursor for peptides such as Substance P), the peptide processing enzyme PAM (peptidylglycine α-amidating monooxygenase), as well as Npy1r, one of the most abundant Npy receptors (Basbaum, Bautista, Scherrer and Julius 2009, Julius and Basbaum 2001). However, genes encoding proteins involved in itch, such as Nppb (brain natriuretic peptide) and Hrh1 (histamine receptor H1) were only expressed at low levels (Usoskin, Furlan, Islam, Abdo, Lonnerberg, Lou, Hjerling-Leffler, Haeggstrom, Kharchenko, Kharchenko, Linnarsson and Ernfors 2015). Similarly, genes responsible for proteins involved in tactile function, including Trpc1 (Transient Receptor Potential Cation Channel Subfamily C Member 1), and those responsible for proprioception, such as Runx3 (Runt related transcription factor 3), exhibited low expression levels. (Le Pichon and Chesler 2014) In contrast, nerve growth factor receptors (Neurotrophic Receptor Tyrosine Kinases 1-3 [Ntrk]) were all expressed at high levels. High expression levels were also observed for transcripts of the two major subpopulations of nociceptors; peptidergic and non-peptidergic (**Fig.4D**).

### Gene expression profiling and enrichment patterns in injured and non-injured cutaneous nociceptors after SCI

To further assess expression profiles of the purified nociceptor population and differences among naïve, sham, and SCI conditions within this nociceptor-enriched population, we focused on expression patterns of gene families that mediate general neuronal functions (Berta, Perrin, Pertin, Tonello, Liu, Chamessian, Kato, Ji and Decosterd 2017, Chiu, Barrett, Williams, Strochlic, Lee, Weyer, Lou, Bryman, Roberson, Ghasemlou, Piccoli, Ahat, Wang, Cobos, Stucky, Ma, Liberles and Woolf 2014). We used differential expression analysis (DESeq2) to analyze significant changes between SCI and naïve or SCI and sham populations (Love, Huber and Anders 2014). Pairwise comparisons of significant genes generated by DESeq2 analysis yielded many differentially expressed genes in each subset (**Suppl.2A-B**). We also assessed differences between sham and naïve populations to exclude significant transcript changes due to laminectomy (**Suppl.2C**).

We focused on expression patterns of gene families which mediate neuronal functions and contribute to pain phenotypes, and found both high expression levels and significant differences within the chloride channel family, Trp channels, glutamate receptors, GABA receptors, potassium channels, sodium channels, and piezo channels (**Fig.5A-G**, p-values in **Table 1**). We also examined ASICs (acid-sensing ion channels), calcium channels, glycine receptors, and P2rx and P2ry families (purinergic receptors), because these are widely studied gene families known to be involved in the development or maintenance of chronic pain (**Suppl.3A-E**). Many channels and receptors were highly expressed within this cell population, but no significant changes among the transcripts were demonstrated for any condition (**Suppl.3**).

**Figure 5.**
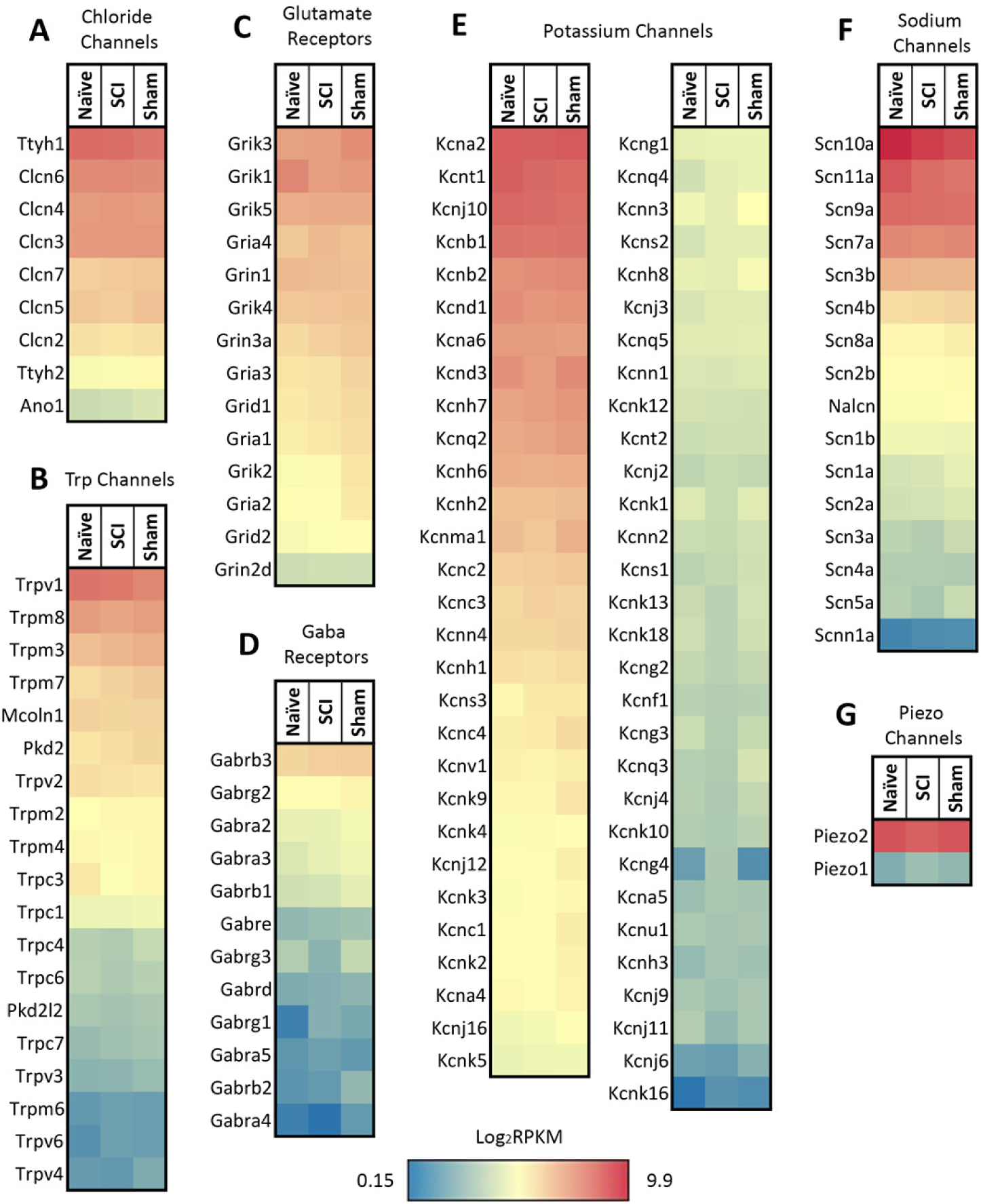
Heatmaps of ion channel transcripts. (A) Chloride channels, (B) TRP channels, (C) glutamate ionotropic receptors, (D) GABA ionotropic receptors, (E) potassium channels, (F) sodium channels, (G) piezo channels. Each family of transcripts includes significant differences in at least one gene 4 days post-SCI. Channels were clustered based on SCI expression level, and graphed by the log_2_ of average RPKM values within each condition. RPKM <1 were not included, N=5 per condition.

**Table 1.**
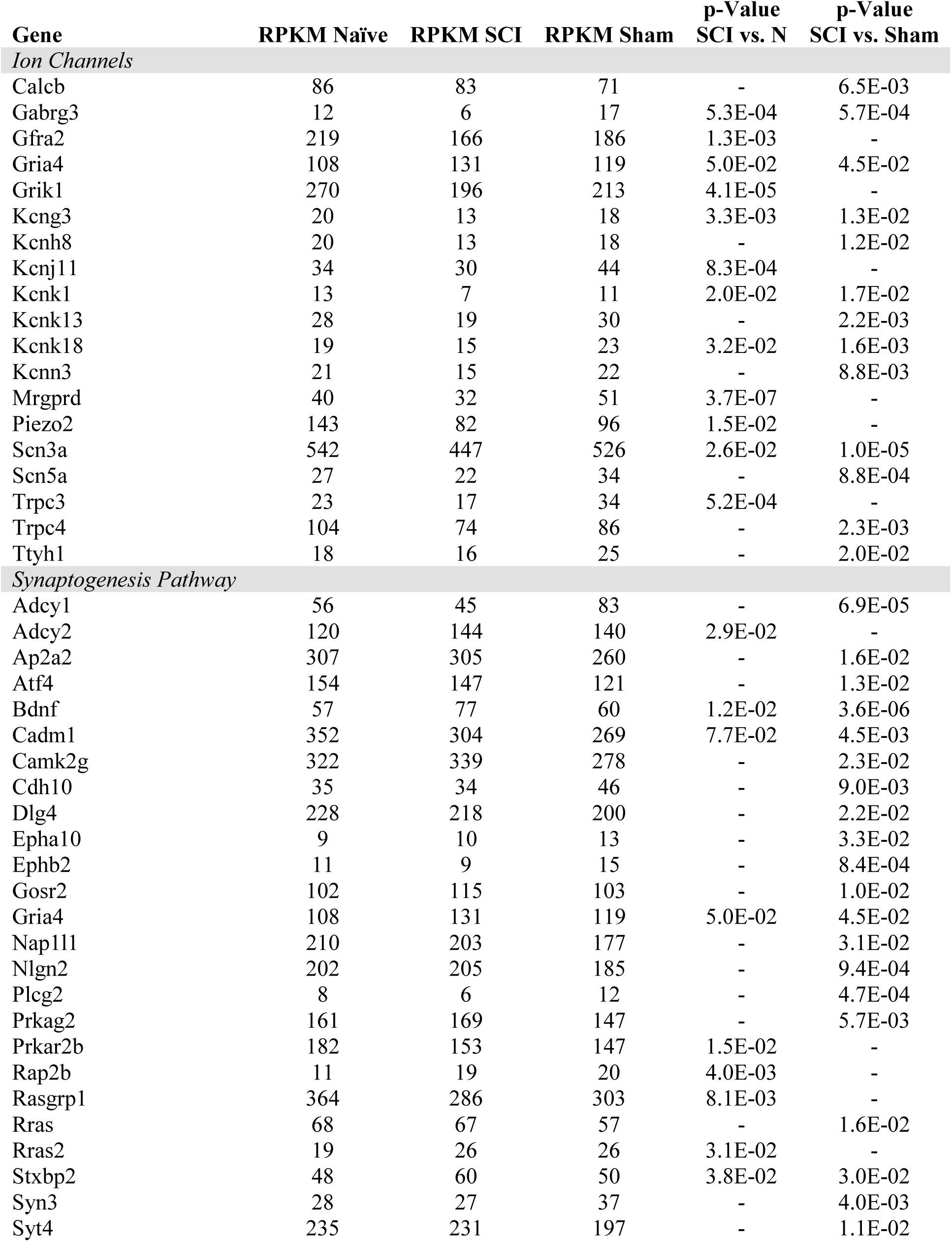
Ion channels and Synaptogenesis. Average RPKM values that significantly differ 4 days post-SCI. DESeq2 p-Value based on SCI vs. naïve or SCI vs. sham comparisons. P-Values that are not listed were >0.05. Transcripts from IPA canonical pathway analysis significantly different 4 days post-SCI. DESeq2 p-Value based on SCI vs. Naïve or SCI vs. Sham comparisons.

Several transcription factors were highly expressed, including Stat3 (signal transducer and activator of transcription 3), Fos (FJB osteosarcoma oncogene), and Jun (Jun Proto-Oncogene) **Suppl.4**). Evidence from various models of neuropathic pain implicates these transcription factors in the development or maintenance of chronic pain (Dominguez et al. 2008, Harris 1998, Naranjo et al. 1991, Tsuda et al. 2011, Xue et al. 2014). Stat3 inhibitors are used to treat peripheral nerve injury-induced hyperexcitability within dorsal horn neurons, pain behaviors, chronic constriction injury, and signaling of IL-6 cytokines (Dominguez, Rivat, Pommier, Mauborgne and Pohl 2008, Tsuda, Kohro, Yano, Tsujikawa, Kitano, Tozaki-Saitoh, Koyanagi, Ohdo, Ji, Salter and Inoue 2011, Xue, Shen, Wang, Hui, Huang and Ma 2014). Previous studies also show that Fos links extracellular events to long-term intracellular changes (such as noxious stimuli) and have established Fos expression as a valid tool to study nociceptive changes (Harris 1998, Naranjo, Mellstrom, Achaval and Sassone-Corsi 1991). Jun also contributes to persistent pain phenotypes following injury (Naranjo, Mellstrom, Achaval and Sassone-Corsi 1991). DESeq2 analysis determined additional transcription factors to be significantly altered by SCI (**Suppl.5**).

### Backlabeled FACS-sorted cutaneous cell transcriptome is distinguished by novel nociceptor-enriched gene patterns

To gain further insight into differentially regulated genes in our isolated population of neurons, we compared our dataset with similar studies on isolated DRGs from publicly available datasets (**Fig. 6**) (Hu et al. 2016, Megat, Ray, Tavares-Ferreira, Moy, Sankaranarayanan, Wanghzou, Fang Lou, Barragan-Iglesias, Campbell, Dussor and Price 2019, Thakur, Crow, Richards, Davey, Levine, Kelleher, Agley, Denk, Harridge and McMahon 2014, Usoskin, Furlan, Islam, Abdo, Lonnerberg, Lou, Hjerling-Leffler, Haeggstrom, Kharchenko, Kharchenko, Linnarsson and Ernfors 2015).). Unsupervised hierarchical clustering of the top 260 genes revealed that a large number of genes display distinct patterns of expression dependent upon the technique used; isolated neurons from all DRG by translating ribosome affinity purification (TRAP) using the Nav1.8^Cre^ mouse (Megat, Ray, Tavares-Ferreira, Moy, Sankaranarayanan, Wanghzou, Fang Lou, Barragan-Iglesias, Campbell, Dussor and Price 2019), single cell isolation from L3-5 DRG (Hu, Huang, Hu, Du, Xue, Zhu and Fan 2016), single cell isolation from L4-L6 DRG (Usoskin, Furlan, Islam, Abdo, Lonnerberg, Lou, Hjerling-Leffler, Haeggstrom, Kharchenko, Kharchenko, Linnarsson and Ernfors 2015), and magnetic cell sorting (MACS) using the Nav1.8 TdTomato mouse (Thakur, Crow, Richards, Davey, Levine, Kelleher, Agley, Denk, Harridge and McMahon 2014) (**Fig.6A**). We found that, while our cutaneous nociceptor-enriched population clustered most closely with the datasets of both TRAP sorted and unsorted DRG from *Megat et. al (Megat, Ray, Tavares-Ferreira, Moy, Sankaranarayanan, Wanghzou, Fang Lou, Barragan-Iglesias, Campbell, Dussor and Price 2019),* our isolated population can be characterized by its own unique data set. Notably, datasets from studies analyzing single cell transcriptomes cluster together, while studies utilizing either TRAP or MACS cluster with their own respective unsorted DRG controls, suggesting that while techniques for isolating specific cell populations can be useful, some variations in gene expression may be attributed to individual differences in cell isolation and RNA extraction methods.

**Figure 6.**
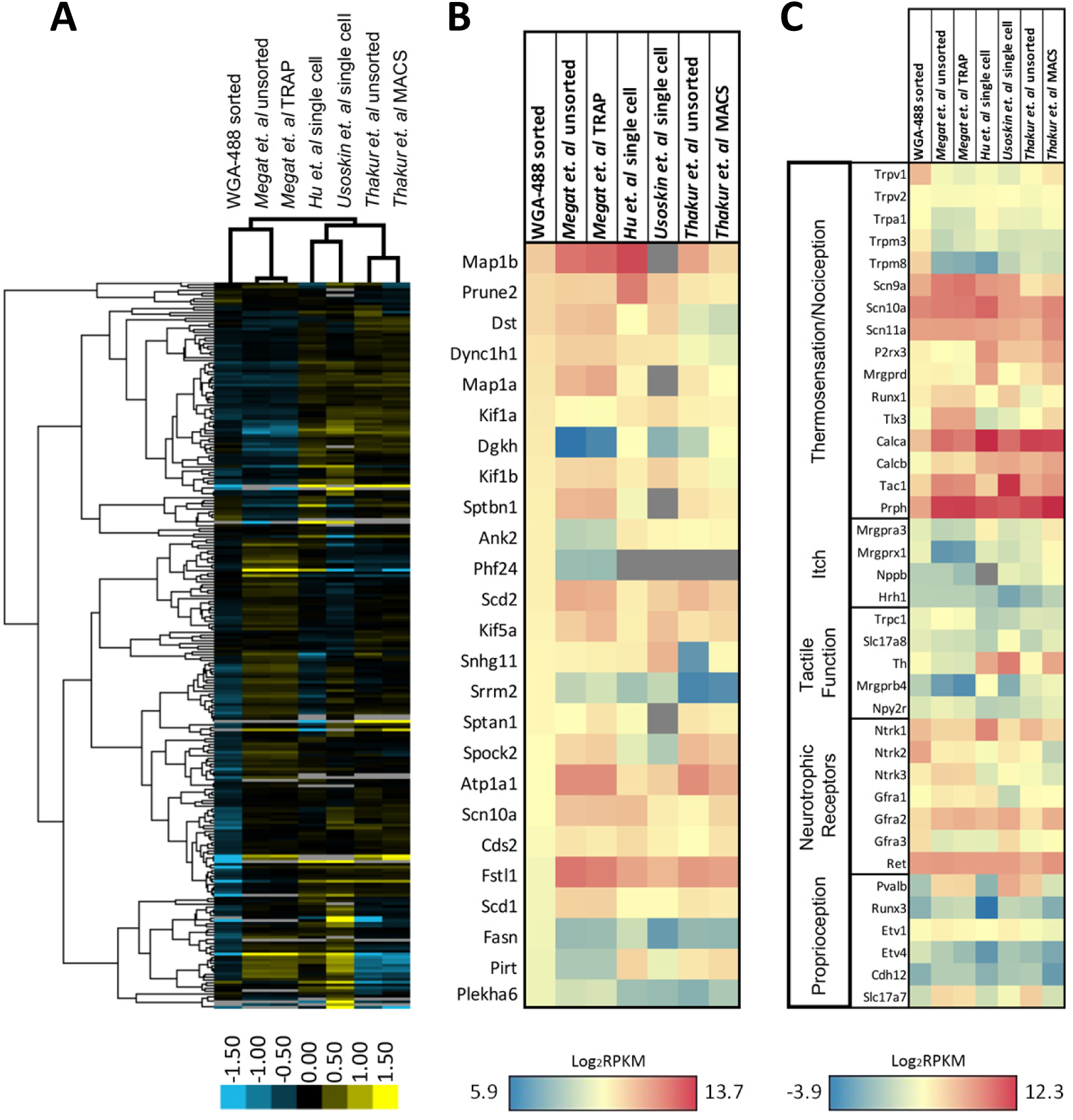
Expression profiling of differentially sorted isolated DRG. (A) Heatmap of the top 260 genes from isolated backlabeled cutaneous neurons of the DRG in comparison to similar studies from publicly available data sets (unsorted dissociated DRG, TRAP-sorted, single cell, or MACS-sorted dissociated DRGs). Comparisons between conditions were made by unsupervised hierarchical clustering. (B) Heatmap of the 25 most significantly enriched genes in our data set in comparison to similar studies from publicly available datasets. Transcripts were graphed by the log_2_ average values within each condition. Datasets that did not express transcripts (RPKM = 0) are depicted in gray. (C) Heatmap of functional somatosensory mediators within the isolated cell population of interest in comparison to similar studies from publicly available data sets. Transcripts were graphed by the log_2_ average values within each condition. Datasets that did not express transcripts (RPKM = 0) are depicted in gray.

To better characterize enriched genes within our population, we graphed the expression of the 25 most significantly enriched genes in our data set in comparison to the same data sets used for hierarchical clustering (**Fig. 6B**). While many of the genes have similarly high expression levels, several genes were unique to our isolated population, including Dgkh (diacylglycerol kinase), Ank2 (ankyrin 2), Phf24 (PHD finger protein 24), Srrm2 (serine/arginine repetitive matrix 2), Fasn (fatty acid synthase), Pirt (phosphoinositide-interacting regulator of transient receptor potential channels), and Plekha6 (pleckstrin homology domain containing, family A member 6). We also compared the expression pattern of our naïve nociceptor-enriched population to the various datasets by again examining known neuronal markers of somatosensation (**Fig. 6C**). As predicted, all of the sorted populations exhibit relatively high expression levels of gene transcripts associated with thermosensation, nociception, or neurotrophic receptors, and comparatively low levels of genes associated with itch, tactile function, or proprioception. It is evident that our cutaneous, nociceptor-enriched population can be defined by its distinct gene expression patterns, in particular high expression levels of the Trp family of genes listed in **Fig. 6C**, as well as Ntrk2, Ntrk3, and Gfra3 neurotrophic receptors.

### Ingenuity pathway analysis (IPA) identified significantly different canonical pathways from cutaneous nociceptors after SCI

Based on DESeq2 analysis, levels of several hundred transcripts in the nociceptor-enriched population were altered by SCI. We restricted IPA input lists to genes that had RPKM values greater than 10 and were statistically different (SCI vs. Naïve or SCI vs. Sham) by DESeq2 analysis (**Fig.7A, B**). Transcripts that exhibited large fold changes included Mrgprb5, Hal, Chrnb4, Cap2, Sez6l, Calb1, Prokr2, Rxfp1, Nxpe2, and Arap3 (**Fig.7A, B**). These genes did not appear in any common significant canonical pathways. IPA identified several pathways that are considered important for inflammatory processes, pain transduction, or the maintenance of chronic pain (**Fig.7C**). This includes calcium signaling, Cxcr4 signaling, neuropathic pain signaling in dorsal horn neurons, opioid signaling, purinergic receptor signaling and synaptic long term potentiation (**Fig.7C**) (Julius and Basbaum 2001, Walters 2012, Walters 2018). The current study focused on significant changes due to SCI pain and not due to post-surgical pain (i.e. changes between sham vs. naïve groups).

**Figure 7.**
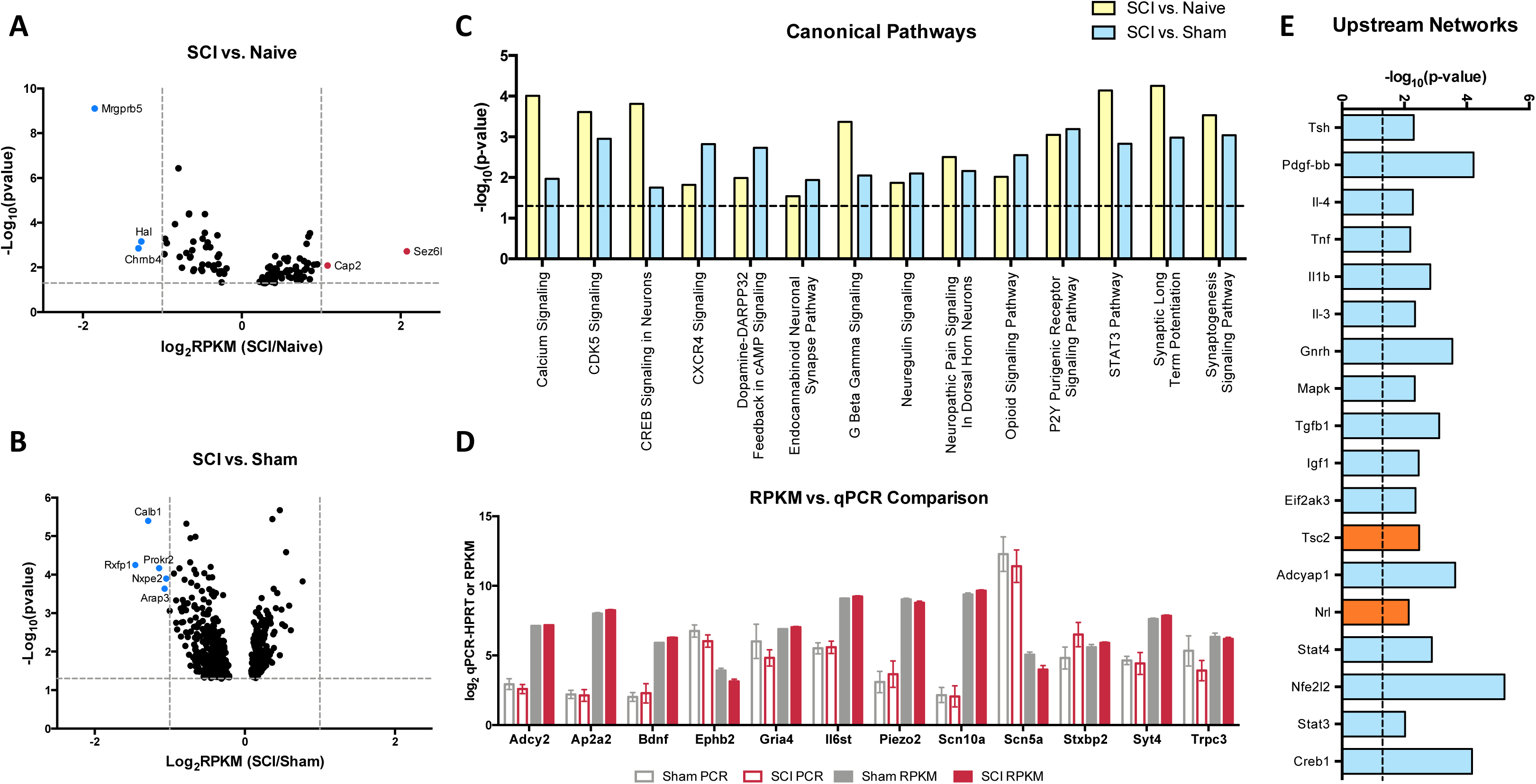
Volcano plots and upstream analysis. (A,B) DESeq2 significant (p>0.05) genes included in IPA analysis comparing SCI vs. naïve or SCI vs. sham conditions. (C) Significant (p<0.05) overlapping canonical pathways predicted by analyzing genes determined by DESeq2 and outlier cutoff in IPA. (D) Comparisons of RPKM values generated by RNAseq and log2 transcript levels (gene-Hprt) validated by qPCR between sham and injured conditions. (E) IPA network analysis predicted networks based on differentially expressed genes between sham and SCI conditions. Network analysis also made predictions about the activation state of the transcript regulator; positive z-score (blue) indicates activation, negative z-score (orange) indicates inhibition.

### Validation of RNASeq data using qPCR

We used qPCR to confirm changes in transcripts of interest from our RNAseq data set. To validate qPCR and RNAseq comparisons, we compared qPCR or RPKM Log_2_ transcript levels of SCI and sham genes of interest (**Fig.7D**). SCI and sham qPCR fold change results were analogous to the RNAseq fold change data set. We focused on the synaptogenesis pathway in particular, as it includes several genes present in overlapping canonical pathways, including changes in receptors involved in organization of excitatory signaling (Ephb) and synapses which may be involved in development of chronic pain (TrkB and BDNF). We validated receptors for significant genes in the synaptogenesis pathway, and genes considered possible contributors to pain that also showed significant differences between conditions Gabrg3, Il6st, Kcng3, Piezo2, Scn5a, Trpc3) (De Jongh et al. 2003, Deng et al. 2018, Devor 2006, Eijkelkamp et al. 2013, Guptarak et al. 2013, Szczot et al. 2018, Waxman et al. 1999, Wickenden 2002, Xia et al. 2015). Additional targets were chosen in order to validate isolation of the correct cell population (Scn10a). Samples for qPCR were collected by backlabeling cutaneous afferents and cell sorting, consistent with samples generated for RNAseq analysis. Following cDNA synthesis, samples were subjected to gene target-specific preamplification using the same primers used for qPCR (**Suppl.6**). Hprt was chosen as the housekeeping gene for qPCR analysis because it was the most constant normalizer transcript from the RPKM data across all 15 mice in the present study (Klenke, Renckhoff, Engler, Peters and Frey 2016, Lima, Gaiteiro, Peixoto, Soares, Neves, Santos and Ferreira 2016).

### IPA network analysis revealed several regulatory interrelationships after SCI

We used IPA upstream network analysis to further interpret the function of the several hundred transcripts significantly altered determined by DESeq2. This method predicted several transcriptional regulators associated with altered expression levels of downstream target genes following SCI **Fig.7E**). The upstream regulator molecules graphed do not show a significant change in RNA expression in response to injury. However, these targets are activated by posttranslational modifications that can alter many of the downstream molecules within its network.

## Discussion

SCI initiates persistent molecular changes in nociceptors, similar to inflammation in models of peripheral injury (Djouhri et al. 2001, Xie et al. 2005). Several studies of SCI pain have evaluated mechanical hypersensitivity from von Frey stimulation above and below the level of thoracic injury, in addition to testing tail withdrawal from heat stimuli (Kramer et al. 2017, Shiao and Lee-Kubli 2018). Additional behavioral studies have demonstrated that SCI animals exhibit significant increases in mechanical and thermal hypersensitivity compared to naïve and sham animals, beginning at 1 month and persisting for several months post-injury (Bedi, Yang, Crook, Du, Wu, Fishman, Grill, Carlton and Walters 2010, Carlton, Du, Tan, Nesic, Hargett, Bopp, Yamani, Lin, Willis and Hulsebosch 2009). However, other studies have asserted that operant behavioral tasks, such as conditioned place preference, are required to effectively study below-level pain in animals (Yezierski 2005). The present study focused on the transition from acute to chronic pain, in the absence of early pain-related behavior, to examine transcriptional differences that occur at much earlier time points rather than a point at which chronic pain is already present. We first tested behavioral differences between naïve and sham mice to identify changes due to the laminectomy, not the spinal cord injury itself, to better determine a time point that captures the transition from acute to chronic pain following SCI. For example, removal of bone and muscle alone could trigger chronic pain-like symptoms, analogous to post-surgical pain reported in humans (Woolf 2011). Surprisingly, there were no significant differences between naïve and sham mice at any time point (**Fig.1A, B**), suggesting that the laminectomy did not produce any locomotor differences or behavioral hypersensitivity 1-7 days post-surgery in mice. By contrast, the spinal cord injury produced clear locomotor differences (**Fig.1A2**).

### Injury and inflammation in SCI

We also considered post-surgical inflammation, using cytokine ELISAs to assess changes in the spinal cord at the level of laminectomy (T8-T11). We wanted to assess changes in cytokine levels for two main reasons; firstly, to confirm that sham mice did not exhibit differences from naïve mice at key timepoints in comparison to SCI mice, and secondly to determine whether sham mice exhibited a prolonged inflammatory response, which could potentially be correlated to the development of chronic pain (Krames 2014). Both pro-inflammatory cytokines TNF-α and IL-1β have been studied in neuroprotection models of SCI. IL-6 has been implicated in neurodegeneration after central nervous system (CNS) injury, and the anti-inflammatory cytokine IL-10 exhibits neuroprotective effects (Donnelly and Popovich 2008, Schmitz and Chew 2008, Zhang et al. 2019). However, we did not find any significant cytokine changes between naïve and sham mice within 7 days of injury, indicating that the laminectomy did not produce a significant inflammatory response at the time points tested. As a positive control for the cytokine ELISAs (Suzuki and Kikkawa 1969), we used the cuprizone model for multiple sclerosis. Key pathological features of the treatment include secretion of proinflammatory cytokines such as TNF-α and IL-1β (Schmitz and Chew 2008). Consistent with previous findings, mice treated with cuprizone exhibited significant increases in TNF-α, IL-1β, and IL-10 relative to naïve mice (**Fig.1C**) (Schmitz and Chew 2008).

Tissue injury can also lead to prolonged functional changes and hyperalgesia that are accompanied by behavioral changes due to increased spontaneous activity of nociceptors (Bedi, Yang, Crook, Du, Wu, Fishman, Grill, Carlton and Walters 2010, Carlton, Du, Tan, Nesic, Hargett, Bopp, Yamani, Lin, Willis and Hulsebosch 2009, Walters 2012). Spontaneous activity in nociceptors following SCI begins at 3 days after injury and persists for at least 8 months (Bedi, Yang, Crook, Du, Wu, Fishman, Grill, Carlton and Walters 2010). This increase in nociceptor activity elicits changes within the spinal dorsal horn, which receives input from these nociceptors, ultimately contributing to spontaneous pain (Dubner and Ruda 1992, Wu et al. 2001). However, the source of hyperexcitability of nociceptors after injury is still unknown. Because spontaneous activity in nociceptors begins by 3 days post-injury, and has been correlated with the generation of persistent pain, we chose to observe transcriptomic changes immediately after this time point at 4 days post-injury (Xie, Strong, Meij, Zhang and Yu 2005). We intentionally excluded large sensory afferents from our experimental model, as the response to SCI by these afferents is transient and has not been directly correlated with pain transduction (Hu, Huang, Hu, Du, Xue, Zhu and Fan 2016, Huang et al. 2006). We focused on an anatomically defined population of nociceptors (projecting from below the level of SCI to hairy hindpaw skin) by back-labeling from peripheral afferent terminals and sorting based on both fluorescence and size. These neurons represent ~10% of dissociated DRG tissue and have a distinct transcriptome (Thakur, Crow, Richards, Davey, Levine, Kelleher, Agley, Denk, Harridge and McMahon 2014). Our approach to identifying anatomically defined small nociceptors is distinct from single cell transcriptome isolation based on the expression of the sodium channel Scn10a (Nav1.8) or expression of advillin (Avil, a marker for all neural crest neurons) (Lopes, Denk and McMahon 2017, Megat, Ray, Tavares-Ferreira, Moy, Sankaranarayanan, Wanghzou, Fang Lou, Barragan-Iglesias, Campbell, Dussor and Price 2019).

### Different DRG neuronal populations in SCI

Different types of sensory neurons are distinct in their responses to injury. It is likely that, even within an identified subpopulation, cells will nonetheless exhibit heterogeneity (Hu, Huang, Hu, Du, Xue, Zhu and Fan 2016). Because injury does not impact all afferents in the same way, we analyzed gene expression changes within the population of nociceptors projecting to the skin below the level of injury. By focusing on small nociceptors innervating dermatomes below the level of injury, we begin to address what unique set of genes within a specific population may be contributing to the burning, stabbing, and shooting pain reported in SCI patients suffering from below-level neuropathic pain (Siddall, McClelland, Rutkowski and Cousins 2003). Our goal was to better understand how SCI affects molecular changes within a specific population of neurons, and how this may contribute to hypersensitivity following SCI. We focused our analysis on sensory neurons from lumbar DRGs (below the level of the SCI) projecting to the hairy hindpaw skin **Fig.2A**). After confirming we had isolated the cell population of interest (**Fig.2B**, **Fig.3**), we used RNAseq to identify changes in gene expression. The use of RNA-Seq has clear advantages over microarrays, since RNA-Seq is not limited to a set of pre-determined transcripts, has a larger dynamic range of transcript expression, and is highly reproducible (Usoskin, Furlan, Islam, Abdo, Lonnerberg, Lou, Hjerling-Leffler, Haeggstrom, Kharchenko, Kharchenko, Linnarsson and Ernfors 2015). By utilizing this technology, we were able to identify transcript changes undetectable with traditional RT-PCR or microarrays (Wang and Zylka 2009). Our RNAseq data from naïve, sham, and injured animals display distinct patterns of somatosensory genes present in this nociceptor-enriched population. In particular, RPKM values show high levels of Scn10a (a marker for nociceptors), purinergic receptor P2rx3, Mrgprd (markers for the non-peptidergic population of nociceptors), and Calca and Calcb (neuropeptide precursors), indicating that we isolated the desired nociceptor specific cell population, and also indicating important genes within the population of sensory neurons projecting to the hairy hindpaw skin (**Fig.4A-D**). Multiple gene transcripts important for itch, tactile function, and proprioception all had relatively low RPKM values, indicating again that we isolated the desired target cell population, and that injury did not induce modifications in the type of stimuli nociceptors transduce (**Fig.4C**). Our population level analysis revealed significant changes after SCI in a number of ion channels and receptors that are already known to play a role in pain or hypersensitivity, such as Piezo2, and transcripts involved in excitatory signaling, such as Grik1 (**Fig.5C**, **Table 1**). However, there were also many genes whose expression and functional roles in persistent pain have yet to be characterized, including Trpc4 and Ttyh1 (**Fig.5A,B**, **Table 1**).

### Transcriptome profiling in studies of DRG

Many studies have utilized RNA-seq technology to gain insight into nociceptor transcriptomes within the DRG and how it changes relative to different pain models (Hu, Huang, Hu, Du, Xue, Zhu and Fan 2016, Megat, Ray, Tavares-Ferreira, Moy, Sankaranarayanan, Wanghzou, Fang Lou, Barragan-Iglesias, Campbell, Dussor and Price 2019, Thakur, Crow, Richards, Davey, Levine, Kelleher, Agley, Denk, Harridge and McMahon 2014, Usoskin, Furlan, Islam, Abdo, Lonnerberg, Lou, Hjerling-Leffler, Haeggstrom, Kharchenko, Kharchenko, Linnarsson and Ernfors 2015). Although previous screens have yielded various nociceptor-specific genes, we have identified a unique pattern of gene expression within a population of nociceptors projecting to the periphery (**Fig. 6**). By cross-comparing our data set with similar studies, we were able to identify our cells of interest and confirm that we obtained cell-type specificity. Notably, many of the current technologies require a transgenic mouse with a cell-type specific reporter. However, by taking advantage of the anatomical organization of the mouse, we were able to use backlabeling and FACS sorting in non-transgenic animals to isolate a nociceptor-enriched population with a low degree of contamination with other cell types. We further implemented this methodology to identify candidate genes in a specific population of neurons below the level of injury in a model of SCI to better understand the heterogenous injury response among the many subtypes of DRG neurons.

Among these comparisons we observed large fold changes in several genes ((**Fig.7A,B**), highlighted in red and blue). However, instead of focusing on larger changes in a small subset of genes with individual functions, we concentrated our analysis on the interaction of many transcripts that were significantly altered after SCI and how these influenced intracellular signaling pathways (**Fig.7C**). Ingenuity Pathway Analysis implicated numerous pathways associated with the progression to persistent pain. We took particular interest in the synaptogenesis signaling pathway as a key player at this 4 day time point, suggesting a role for synaptic plasticity in the transition from acute to chronic pain after SCI. In addition to its relevance within our model, many of the transcripts involved in synaptic plasticity overlapped with several other pathways (**Fig.7C**) and had RPKM values that could be validated by qPCR (**Fig.7D**). Synaptogenesis is typically associated with developmental processes, including axon guidance and synapse formation (Klein 2004). However, activation of various signaling pathways involved in synaptogenesis may also contribute to pain; for example persistent pain is supported via changes in synaptic signaling, neuronal plasticity, and long term potentiation, and may form memory-like networks for painful signals that allow persistent pain to occur long after the initial injury (Khangura et al. 2019, Kobayashi et al. 2007).

Included in the many of the genes of interest within the synaptogenesis pathway, Ephb2 was significantly down regulated post-injury (**Table 1**). The gene transcript is part of the Ephrin tyrosine kinase receptor protein family that is expressed in laminae I-III of the spinal dorsal horn on small and medium sized DRG neurons (presumably nociceptors) (Bundesen et al. 2003). Ephb receptors regulate synaptic activity in the spinal cord and contribute to persistent pain associated with NMDA activity (Khangura, Sharma, Bali, Singh and Jaggi 2019). Numerous receptor tyrosine kinases localize to synapses and contribute to synaptogenesis in addition to EphB receptors, including Trk receptors (Biederer and Stagi 2008). Ntrk2, the receptor for BDNF, was highly expressed in this cell population; commensurately, BDNF transcript levels were significantly upregulated in the injured population (**Table 1**). Camk2g transcript levels were significantly increased in the SCI population of cutaneous nociceptors as well, and recent work has shown phosphorylation of Camk2g induces Bdnf mRNA transcription (Yan et al. 2016). This parallels increasing evidence that neuronal activity (such as increased activity or hyperexcitability) activates alternative neuronal circuits through activity-regulated genes, such as *BDNF* (Lu et al. 2009). Many of the genes significantly altered in the synaptogenesis pathway may function together to generate neuropathic pain (**Suppl.7**).

### Networks of genes coordinating responses to SCI

Regulatory interrelationships predicted by the IPA program were also examined (**Fig.7E**). Many of these networks are related to inflammatory signaling mechanisms, suggesting a link between pro-inflammatory signaling and synaptic transmission (Medelin et al. 2018). Previous work has associated inflammatory mechanisms to diseases of the CNS, including multiple sclerosis, Alzheimer’s disease, and Parkinson’s disease (Medelin, Giacco, Aldinucci, Castronovo, Bonechi, Sibilla, Tanturli, Torcia, Ballerini, Cozzolino and Ballerini 2018). The mechanisms through which inflammation prompts changes in synaptic transmission are not fully understood, but several of the classic pro-inflammatory cytokines (including TNF-α, IL-6, IL-1β) have been shown to contribute to a decrease in hippocampal neurogenesis, and could be playing a similar role within the spinal cord following injury (Kohman and Rhodes 2013). Overall, pro-inflammatory conditions have been associated with increases in post-synaptic NMDA and AMPA receptors, and inhibition of GABAergic receptors (Medelin, Giacco, Aldinucci, Castronovo, Bonechi, Sibilla, Tanturli, Torcia, Ballerini, Cozzolino and Ballerini 2018).

## Conclusion

Molecular changes typically reflect phenotypic characteristics, and our data show changes in gene expression 4 days after injury, suggesting that many of these genes may be responsible for the development of spontaneous activity reported elsewhere (Bedi, Yang, Crook, Du, Wu, Fishman, Grill, Carlton and Walters 2010, Yang, Wu, Hadden, Odem, Zuo, Crook, Frost and Walters 2014). We recognize that RNA-Seq of batched neurons elucidated changes in gene targets in a subpopulation of cells, but averaging occurred when pooling large numbers of cells, precluding analysis at the level of the single cell (Haque et al. 2017). Further analysis at the single cell level of cutaneous nociceptors will clarify the contributions of specific subpopulations (non-peptidergic versus peptidergic) to chronic pain after SCI. Functional studies are also needed to analyze the roles of this specific cell population, to better understand the connectivity and plasticity of the CNS and PNS. While our results begin to address prospective gene networks that may contribute to the development of chronic pain, additional behavioral testing in conjunction with targeting of specific biological mechanisms until the chronic pain phase is necessary in order to attribute specific transcriptional changes to pain phenotypes. The DRG nociceptor preparation isolated by backlabeling with WGA-488 and FACS has many applications in molecular studies. We have demonstrated how the cutaneous nociceptor transcriptome is altered following SCI to gain novel biological insight into disease mechanisms in a cell-type specific approach. It is evident that the transition from acute to chronic pain occurs in distinct steps that involve numerous signaling pathways, providing a host of potential new drug targets.

## Author Contributions

J.Y. and R.M. designed research; J.Y. performed research; J.Y. and R.M. analyzed data; J.Y., I.M. and R.M. wrote the paper.

## Acknowledgments

We thank Dr. Nicholas Wasko and Dr. Robert Clarke for providing us with cuprizone treated mice, and Dr. Evan Jellison for his help and expertise with flow cytometry. We acknowledge Dr. Bo Reese and the Center for Genome Innovation, Institute for Systems Genomics, University of Connecticut for library construction and RNA sequencing services. We also acknowledge Dr. Vijender Singh and the Computational Biology Core, Institute for Systems Genomics, University of Connecticut for base-calling, read alignment, and assembly of individual transcripts to align to the genome. This work was supported by NIH DK032948 (REM) and the University of Connecticut Graduate School.

## Data Availability

The datasets generated during and/ or analyzed during the current study are available in Gene Expression Omnibus repository under the series record number GSE132552, all other data generated during this study are included in this published article (and its Supplementary Information files).

---

**Supplementary Figure 1.**
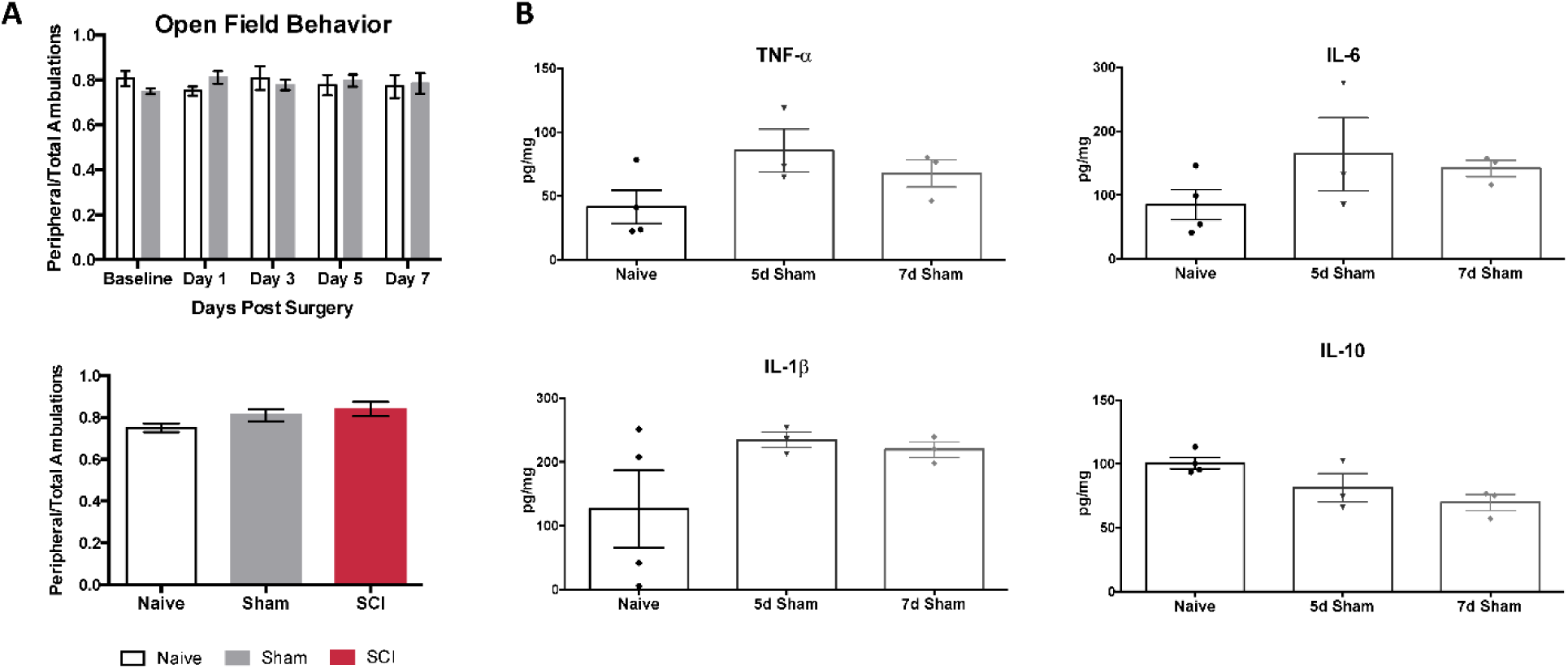
Mobility, cytokine controls. (A) Open field behavior (10-minute trials) conducted on naïve and sham mice 0,1,3,5 and 7 days post-surgery does not differ significantly at any time point in time spent in periphery, N=6 each. Testing done 1 day post-SCI also does not differ significantly in time spent in periphery, N=6 naïve, N=6 sham, N=4 SCI. (B) Cytokine ELISAs on spinal cord segments at the level of laminectomy (T8-T11) show no significant differences between naïve and sham mice 5 or 7 days post-surgery.

**Supplementary Figure 2.**
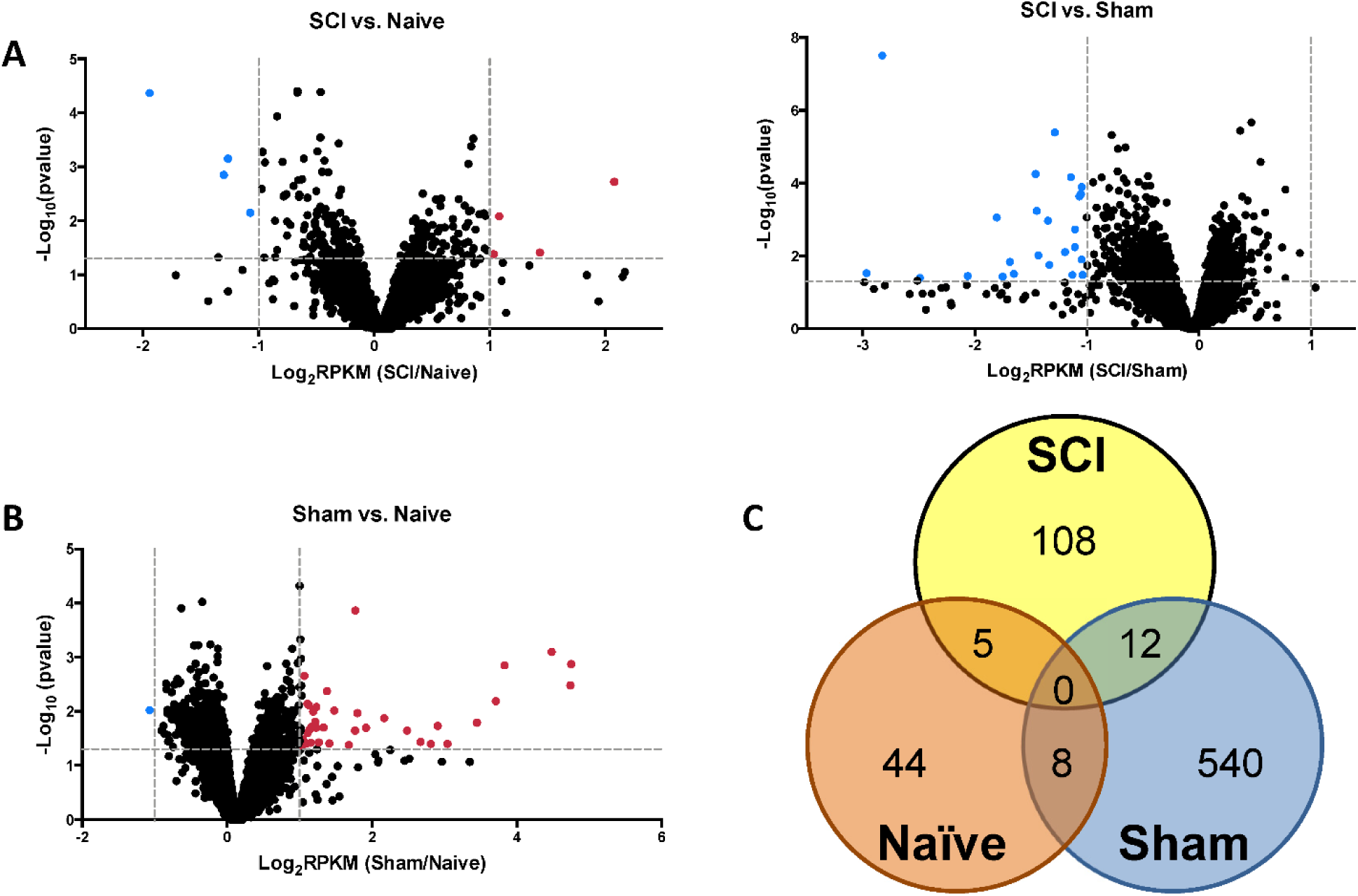
Volcano plots, Venn diagram. (A) Volcano plot of RNAseq transcript p-Values calculated by DESeq2 comparing SCI vs. naïve, SCI vs. sham, or (B) sham vs. naïve conditions, RPKM >10. (C) Venn diagram of statistically significant genes from the RNAseq data set determined by an overlap of DESeq2 significant genes (p<0.05) and outlier removal, with a cutoff excluding RPKM values <10.

**Supplementary Figure 3.**
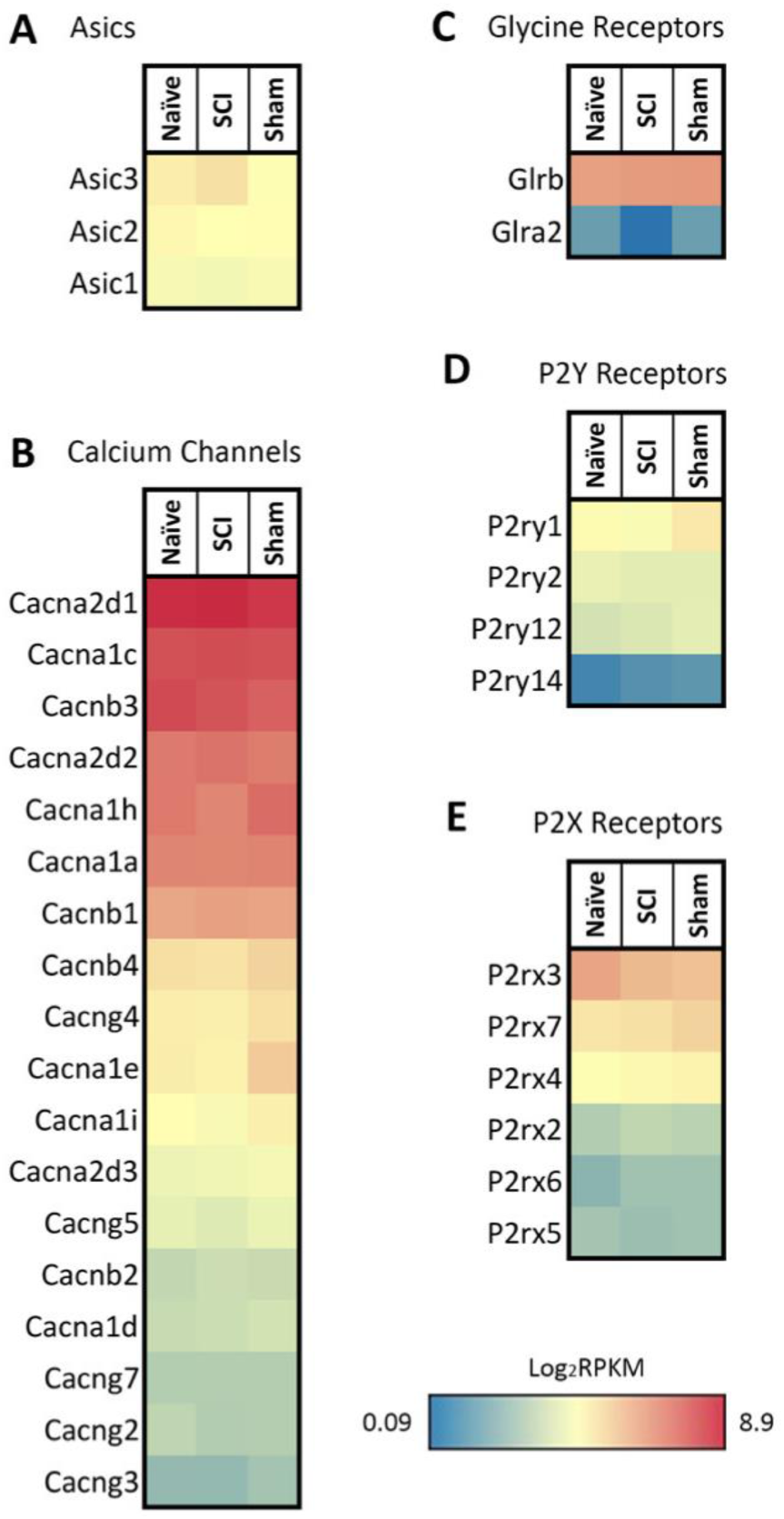
Ion channel heatmaps. Acid sensing ion channels (Asics), calcium channels, glycine receptors, and purinergic receptors (P2Y, P2X). Expression patterns are similar across all three conditions. Despite their known relevance in pain transduction, no significant changes were observed at the 4 day time point tested. RPKM <1 were not included.

**Supplementary Figure 4.**
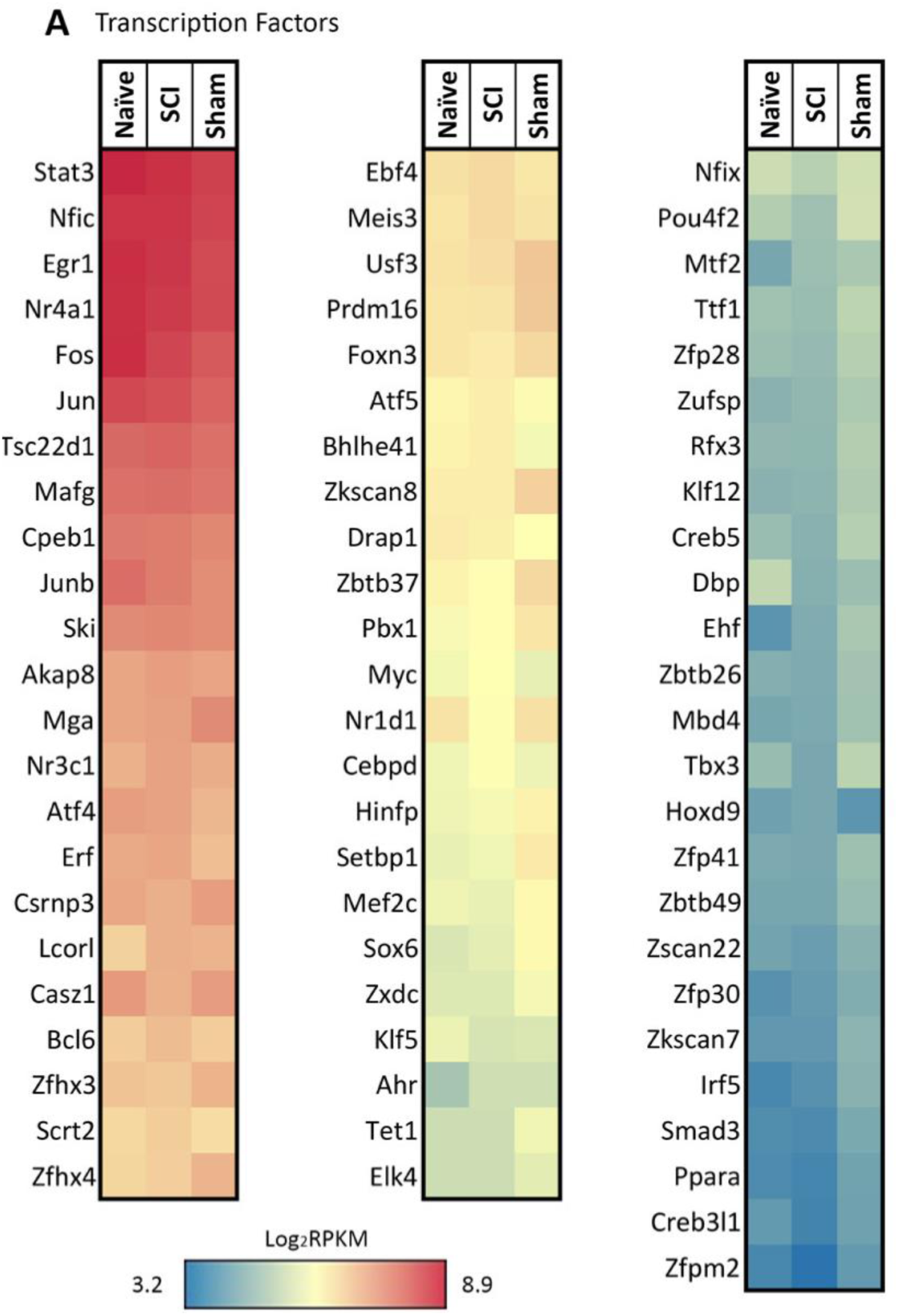
Transcription factor heatmap. Significant changes between SCI vs. naïve or SCI vs. sham conditions by DESeq2: Ahr, Atf4, Cpeb1, Creb3l1, Csrnp3, Drap1, Egr1, Erf, Foxn3, Irf5, Jun, Junb, Mafg, Mef2c, Meis3, Myc, Nr3c1, Nr4a1, Pbx1, Tbx3, Tet1, Zfhx3, Zfp28, Zfp30, Zfp41, Zkscan8, Zscan22. RPKM <1 were not included.

**Supplementary Figure 5.**
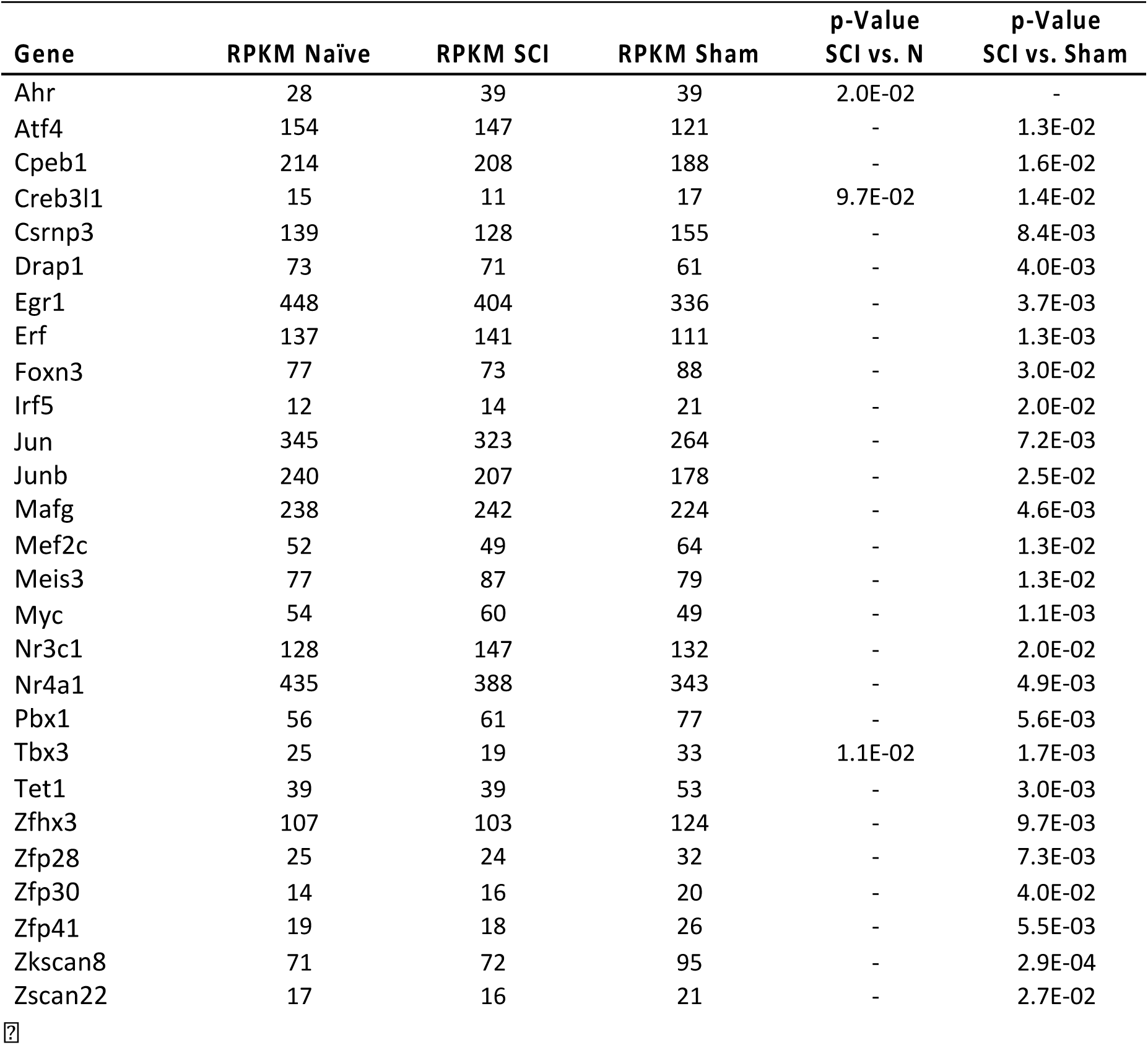
Transcription factors. Transcript levels that significantly differ 4 days post-SCI. DESeq2 p-Value based on SCI vs Naïve or SCI vs Sham comparisons. P-Values that are not listed were >0.05.

**Supplementary Figure 6.**
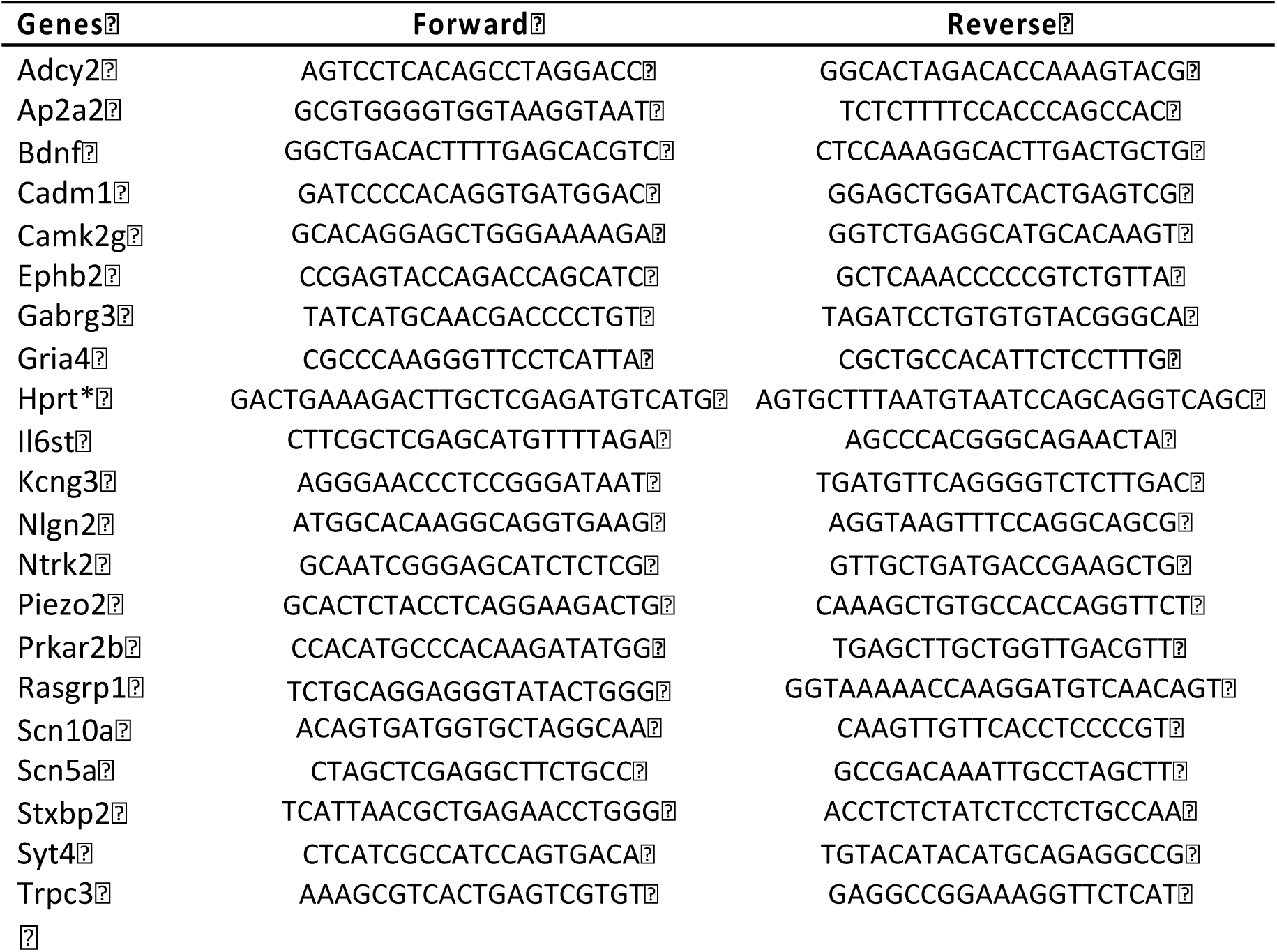
Primer list. Primers for voltage-gated channels, receptors, Trp channels, or involvement in the synaptogenesis pathway were designed for PCR products 111-143 bp in size, Tm=59.5-63.5C, validated on whole DRG tissue before preamplification. Hprt was the least variable gene based on RNAseq results.

**Supplementary Figure 7.**
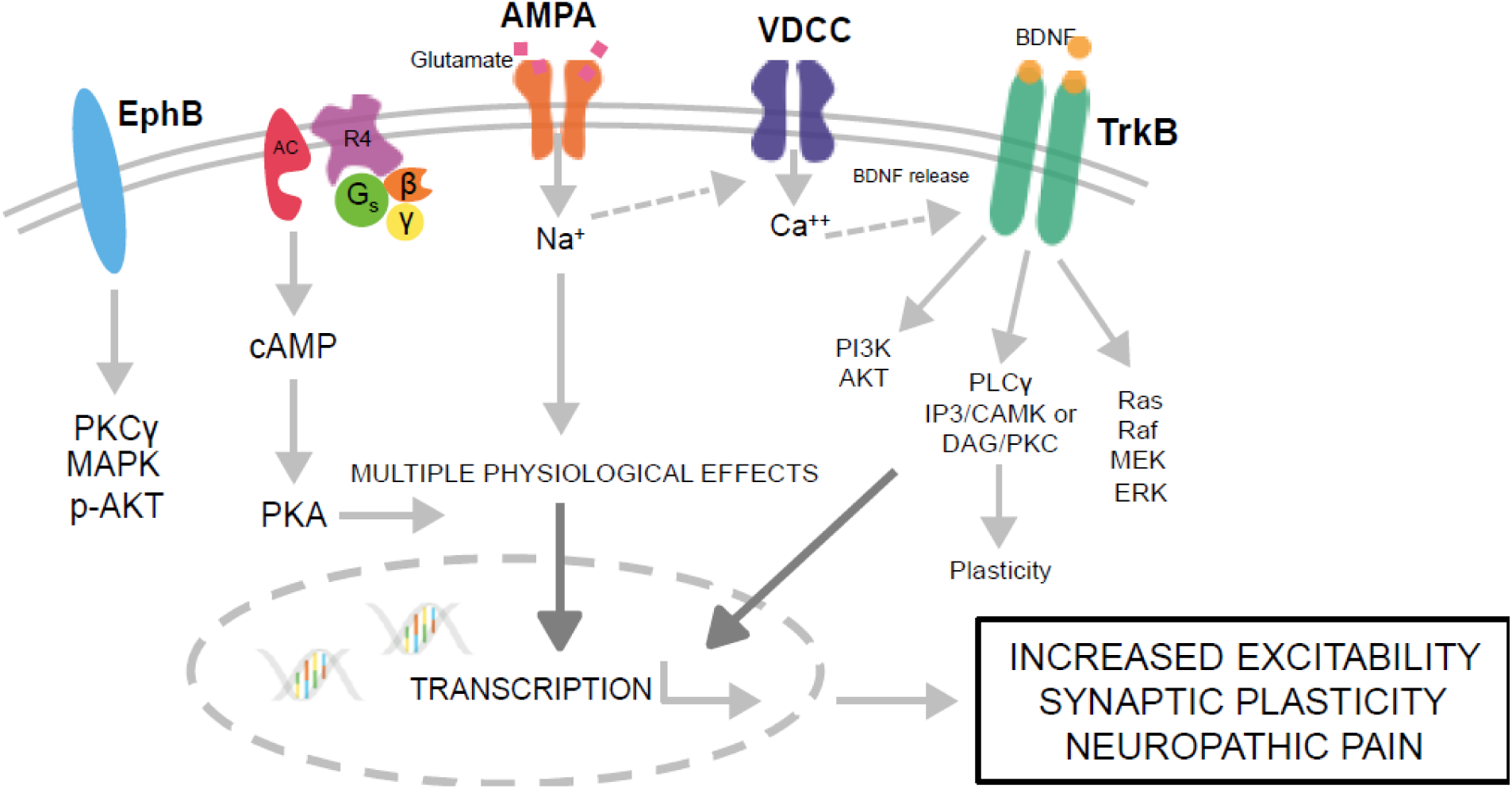
Proposed model of how this signaling pathway may be contributing to the onset of chronic pain 4 days post-SCI in DRG distal to the site of injury. RNAseq data, IPA analysis, and qPCR validation suggest Ntrk2 (TrkB) signaling may play a role during the transition from acute to chronic pain at 4 days post-SCI.

